# Cholinergic circuit genes in the healthy brain are differentially expressed in regions that exhibit gray matter loss in Parkinson’s disease

**DOI:** 10.1101/2019.12.17.875880

**Authors:** Arlin Keo, Oleh Dzyubachyk, Jeroen van der Grond, Anne Hafkemeijer, Wilma D.J. van de Berg, Jacobus J. van Hilten, Marcel J. T. Reinders, Ahmed Mahfouz

## Abstract

Structural covariance networks are able to identify functionally organized brain regions by gray matter volume covariance. In Parkinson’s disease, the posterior cingulate network and anterior cingulate network showed decreased gray matter and therefore we examined the underlying molecular processes of these anatomical networks in the healthy brain. Whole brain transcriptomics from post-mortem samples from healthy adults, revealed upregulation of genes associated with serotonin, GPCR, GABA, glutamate, and RAS signaling pathways in these PD-related regions. Our results also suggest involvement of the cholinergic circuit, in which genes *NPPA, SOSTDC1*, and *TYRP1* may play a protective role. Furthermore, both networks were associated with memory and neuropsychiatric disorders that overlap with Parkinson’s disease symptoms. The identified genes and pathways contribute to healthy functions of the posterior and anterior cingulate networks and disruptions to these functions may in turn contribute to the pathological and clinical events observed in Parkinson’s disease.

## Introduction

Parkinson’s disease (PD) is a neurodegenerative disorder pathologically defined by the loss of dopaminergic neurons in the substantia nigra and the presence of Lewy bodies in selective brain regions (Goedert et al. 2012). Clinical symptoms involve impairment of diverse motor and non-motor symptoms that get progressively worse over time. The decline in clinical performance has been associated with changes in morphological properties of structural and functional neuroimaging networks (Lucas-Jiménez et al. 2016; Wang et al. 2016; de Schipper et al. 2017). In turn, studies have investigated the relationship between these imaging networks and genetic variants in PD-related genes to provide new insights into the pathogenesis of PD (van der Vegt et al. 2009). However, less is known about the functional pathways that underlie the spatial organization of brain regions contributing to PD. To identify the molecular mechanisms underlying changes in structural and functional networks in PD, imaging data has been integrated with brain wide expression atlases of the healthy brain (Hawrylycz et al. 2015; Arnatkevičiūtė et al. 2019). Regional brain atrophy in PD patients was found to be correlated with the expression of genes implicated in trans-synaptic alpha-synuclein transfer (Freeze et al. 2018) and a loss of regional connectivity in PD patients was correlated with the regional expression of *MAPT* in the healthy brain (Rittman et al. 2016). These studies show that combining imaging data in PD and gene expression from the healthy brain can shed light on the molecular mechanisms underlying the observed differences between PD and controls.

Structural covariance networks (SCNs) can reveal functional network organizations by identifying brain regions that co-vary in gray matter volume across a population (Alexander-Bloch, Giedd, et al. 2013). SCNs have been shown to be dysregulated in different neurological disorders (Alexander-Bloch, Raznahan, et al. 2013; Spreng and Turner 2013; Coppen et al. 2016; Huang et al. 2017; Liu et al. 2019), and variations in SCNs can be explained by transcriptomic similarity and structural connectivity (Romero-Garcia et al. 2018; Yee et al. 2018). Hafkemeijer et al. (Hafkemeijer et al. 2014) identified nine SCNs based on gray matter variation among healthy middle-aged to older adults. Gray matter volume in four of these nine networks was negatively associated with age: a subcortical network, sensorimotor network, posterior cingulate networks, and anterior cingulate network. Two of these networks were found to show loss of gray matter volume in PD patients beyond the effects of aging: the posterior cingulate network and anterior cingulate network (de Schipper et al. 2017). These two networks were negatively associated with cognitive impairment and daytime sleepiness, respectively. Yet, it is unclear which molecular mechanisms contribute to these differences in SCNs in individuals with PD.

In this study, we aim to identify the molecular mechanisms within the healthy brain underlying the loss of integrity and atrophy in the anterior and posterior cingulate networks in PD patients. By integrating the nine SCNs with spatial gene expression data from the Allen Human Brain Atlas, we investigate whether expression patterns in the normal healthy brain are associated with patterns of gray matter loss in PD. We show that genes highly expressed in the posterior and anterior cingulate networks were associated with multiple neurotransmitter signaling pathways and involved in memory-related, pain-related, and neuropsychiatric disorders. In addition, both anatomical networks showed high expression of cholinergic gene markers known to act as regulators of extracellular signaling. Our results provide new insights into the molecular processes underlying anatomical network function and aids in understanding the selective progression of PD.

## Materials and Methods

### Transcriptomic data preprocessing

To understand transcriptomic signatures of nine anatomical networks of the healthy brain, we used normalized gene expression data from the Allen Human Brain Atlas (AHBA), a post-mortem microarray data set of 3,702 anatomical brain regions from six healthy individuals (5 males and 1 female, mean age 42, range 24–57 years) (Hawrylycz et al. 2015). We downloaded the data from http://human.brain-map.org/. To filter and map probes to genes, the data was concatenated across the six donors. We removed 10,521 probes with missing Entrez IDs, and 6,068 probes with low presence as they were expressed above background in <1% of samples (PA-call containing presence/absence flag) (Hawrylycz et al. 2015). The remaining 44,072 probes were mapped to 20,017 genes with unique Entrez IDs using the *collapseRows*-function in R-package WGCNA v1.64.1 (Langfelder and Horvath 2008) as follows: i) if there is one probe, that one probe is chosen, ii) if there are two probes, the one with maximum variance across all samples is chosen (method=”maxRowVariance”), iii) if there are more than two probes, the probe with the highest connectivity (summed adjacency) is chosen (connectivityBasedCollapsing=TRUE).

For visualization of gene expression in heatmaps, data was Z-score normalized across all samples for each brain donor separately. Heatmaps were plotted using R-package ComplexHeatmap v2.0.0 (Gu et al. 2016). Genes were clustered using complete linkage with Euclidean distances. The same color scale was used for all heatmaps.

### Mapping AHBA samples to SCNs of the healthy brain

We focused on anatomical networks that were previously defined based on whole brain gray matter volume covariation in 370 middle-aged to older adults between 45 and 85 years; for more detailed information on the networks see Hafkemeijer et al. 2014. Nine networks were defined and named according to the presence of the main structures: thalamus (network A), lateral occipital cortex (network B), posterior cingulate cortex (network C), anterior cingulate cortex (network D), temporal pole (network E), putamen (network F), and cerebellum (networks G, H, and I). The same networks were previously investigated for loss of integrity in 159 PD patients from the same age range, where the posterior cingulate network (C) and anterior cingulate network (D) showed decreased gray matter; for demographic and clinical information see de Schipper et al. 2017. All samples from each one of the six donors in AHBA were mapped to regions defined by the nine SCNs in MNI coordinate space. The mapping divided all the samples into two sets depending on their position inside (1) or outside (0) the SCN mask.

### Differential expression analysis

Differential expression analysis was performed between each of the two PD-related networks (posterior and anterior cingulate cortex) and the other 7 non-PD-related networks together. A two-tailed t-test was used for each gene and the analysis was done separately for each donor from AHBA. Since the microarray data was log^2^-transformed, the mean expression difference is interpreted as the log^2^-transformed fold-change (FC). The effect sizes for each one of the six donors were combined by meta-analysis (metafor R-package 2.0). For the meta-analysis, a random effects model was applied which assumes that each brain is considered to be from a larger population of brains and therefore takes the^2^ within-brain and between-brain variance into account. The between-brain variance (tau) was estimated with the Dersimonian-Delaird model. Variances and confidence intervals were obtained using the escalc-function. The significance of summary effect sizes was assessed through a two-sided t-test (H^0^: FC=0; unequal variances). *P*-values of the effect sizes were Benjamini-Hochberg (BH) corrected for all 20,017 genes.

### Pathway analysis

Pathway analysis was done with the ReactomePA R-package version 1.28 using the function *enrichPathway* searching for human pathways. All 20,017 genes in the AHBA dataset were set as background genes. Pathways with a minimum size of 10 genes were significant when BH-corrected *P* < 0.05.

### Cell-type marker enrichment

Gene markers for 28 cell-types were downloaded from the Neuroexpresso database (http://neuroexpresso.org/) using markers from all brain regions. These have been identified in a cross-laboratory dataset of cell-type specific transcriptomes from the mouse brain (Mancarci et al. 2017). To assess their expression, Entrez IDs of the mouse cell-type specific markers were converted to human homologs (homologene R-package version 1.4) and filtered for genes present in the AHBA dataset (Supplementary Table 1). Two markers with different mouse gene IDs (14972, *H2-K1*, microglial, and 15006, *H2-Q1* serotonergic), were converted to the same human gene ID (3105, *HLA-A*), and therefore removed before analysis. For cell-type enrichment, we assessed which cell-type markers were overrepresented among the differentially expressed genes. For 17 cell-types that had at least six markers (astrocyte, Bergmann, cerebellar granule, dentate granule, ependymal, GabaReln, hypocretinergic, microglia, activated microglia, deactivated microglia, noradrenergic, oligo, purkinje, serotonergic, spinal cord cholinergic, spiny, and thalamus cholinergic), we assessed the significance with the hypergeometric test and *P*-values were corrected for all 17 cell-types (BH-corrected *P* < 0.05).

### Enrichment of disease-associated genes

Differentially expressed genes were also assessed for overrepresentation of disease-associated genes from DisGeNET (Piñero et al. 2017). A table of 628,685 gene-disease associations were obtained from DisGeNET version 6.0 (July, 2019) from http://www.disgenet.org/. A hypergeometric test was used to assess the significance of overlapping genes (P < 0.05), and *P*-values were BH-corrected for 24,166 diseases. The odds ratio (OR) for cell-type and disease enrichment was calculated using the DescTools R-package.

## Results

### Transcriptomics of the posterior and anterior cingulate networks

We analyzed the transcriptomes of healthy subjects across nine anatomical networks defined by structural covariance of gray matter volume among healthy middle-aged to older adults (Hafkemeijer et al. 2014). We focused on the posterior cingulate network and the anterior cingulate network that showed loss of gray matter in PD patients which seemed to be associated with cognition and excessive daytime sleepiness (Figure 1) (de Schipper et al. 2017). For this we used the AHBA microarray dataset of spatial gene expression in post-mortem brains. AHBA samples were mapped to each one of the nine networks A-I (Table 1). We characterized the transcriptional signature of the two PD-related networks (Network C and Network D) in these AHBA samples by comparing their gene expression pattern to the remaining seven networks together (non-PD-related).

**Figure 1.**
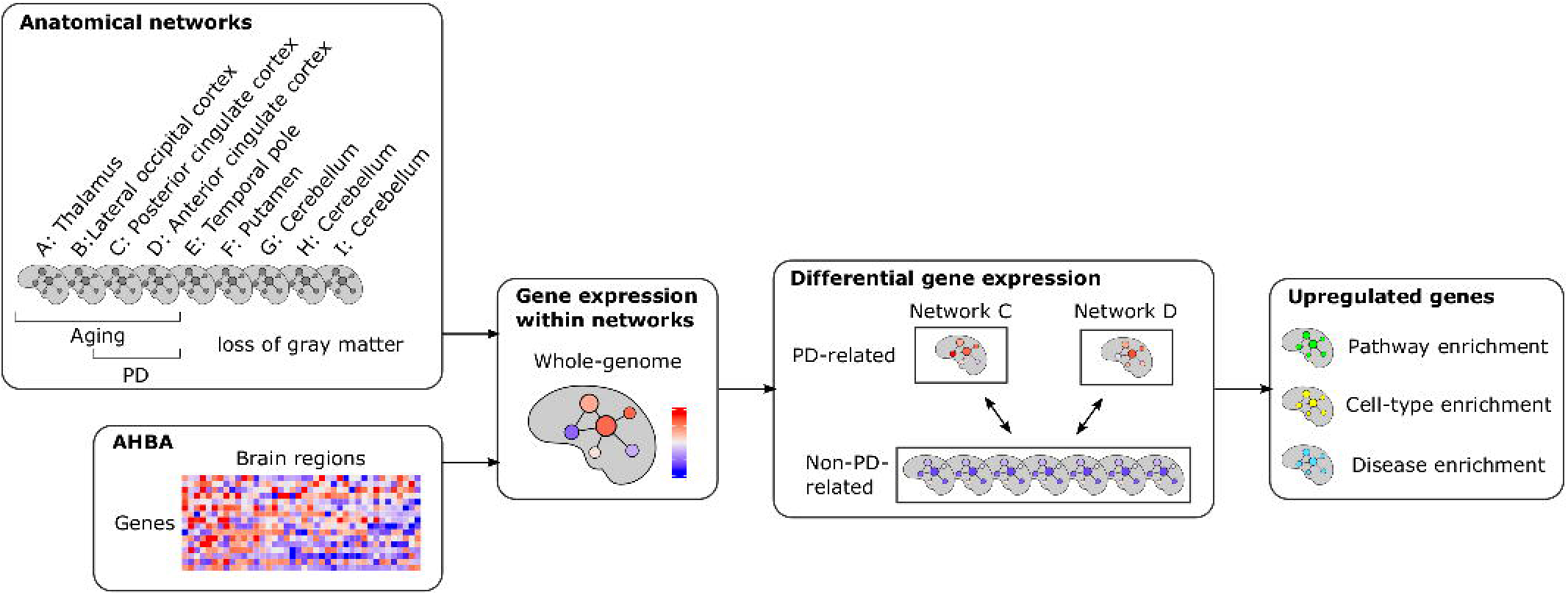
Study overview. Transcriptomic data from AHBA were mapped to nine anatomical networks that have been defined based on healthy subjects. The posterior cingulate network and anterior cingulate network have been associated with gray matter loss in PD (PD-related), while the seven remaining networks were not related to PD (non-PD-related). We compared gene expression in each of the two PD-related networks to gene expression in all seven non-PD-related networks. Genes that were upregulated in the two PD-related networks were assessed for the overrepresentation of pathway-specific genes, cell-type marker genes, and disease-associated genes.

**Table 1.** Number of samples from AHBA within SCN networks.

| Donors | Network |  |  |  |  |  |  |  |  |
| --- | --- | --- | --- | --- | --- | --- | --- | --- | --- |
|  | A | B | C | D | E | F | G | H | I |
| Donor 9861 | 72 | 67 | 157 | 47 | 74 | 90 | 26 | 39 | 83 |
| Donor 10021 | 79 | 46 | 121 | 65 | 49 | 84 | 25 | 55 | 91 |
| Donor 12876 | 37 | 24 | 57 | 28 | 42 | 45 | 6 | 17 | 25 |
| Donor 14380 | 38 | 33 | 52 | 30 | 45 | 61 | 7 | 27 | 53 |
| Donor 15496 | 34 | 24 | 41 | 21 | 39 | 55 | 13 | 24 | 69 |
| Donor 15697 | 49 | 20 | 38 | 33 | 47 | 64 | 29 | 37 | 49 |
| Total | 309 | 214 | 466 | 224 | 296 | 399 | 106 | 199 | 370 |

A: Thalamus; B: Lateral occipital cortex, C: Posterior cingulate cortex, D: Anterior cingulate cortex, E: Temporal pole; F: Putamen; G, H, I: Cerebellum.

Genes were differentially expressed within the posterior cingulate network or the anterior cingulate network compared to the other networks combined when absolute fold-change (FC) > 1 and Benjamini-Hochberg (BH) corrected *P*-value < 0.05. Differential expression analysis showed a large overlap of genes that were differentially expressed in the same direction in the two networks. We found that 73 genes in the posterior cingulate network and 39 genes in anterior cingulate network were downregulated, of which 25 genes overlapped between both networks (Figure 2A and B, Supplementary Table 2 and 3). Furthermore, 200 genes in the posterior cingulate network and 269 genes in anterior cingulate network were upregulated, for which 144 genes overlapped (Supplementary Table 4 and 5). Among the differentially expressed genes in the posterior and anterior cingulate networks, we further assessed the presence of PD-implicated genes identified in familial and genome-wide association studies (Bonifati 2014; Nalls et al. 2014; Chang et al. 2017). Although mutations in these genes may impact disease outcome, none of these genes were differentially expressed in the posterior cingulate network and anterior cingulate network.

**Figure 2.**
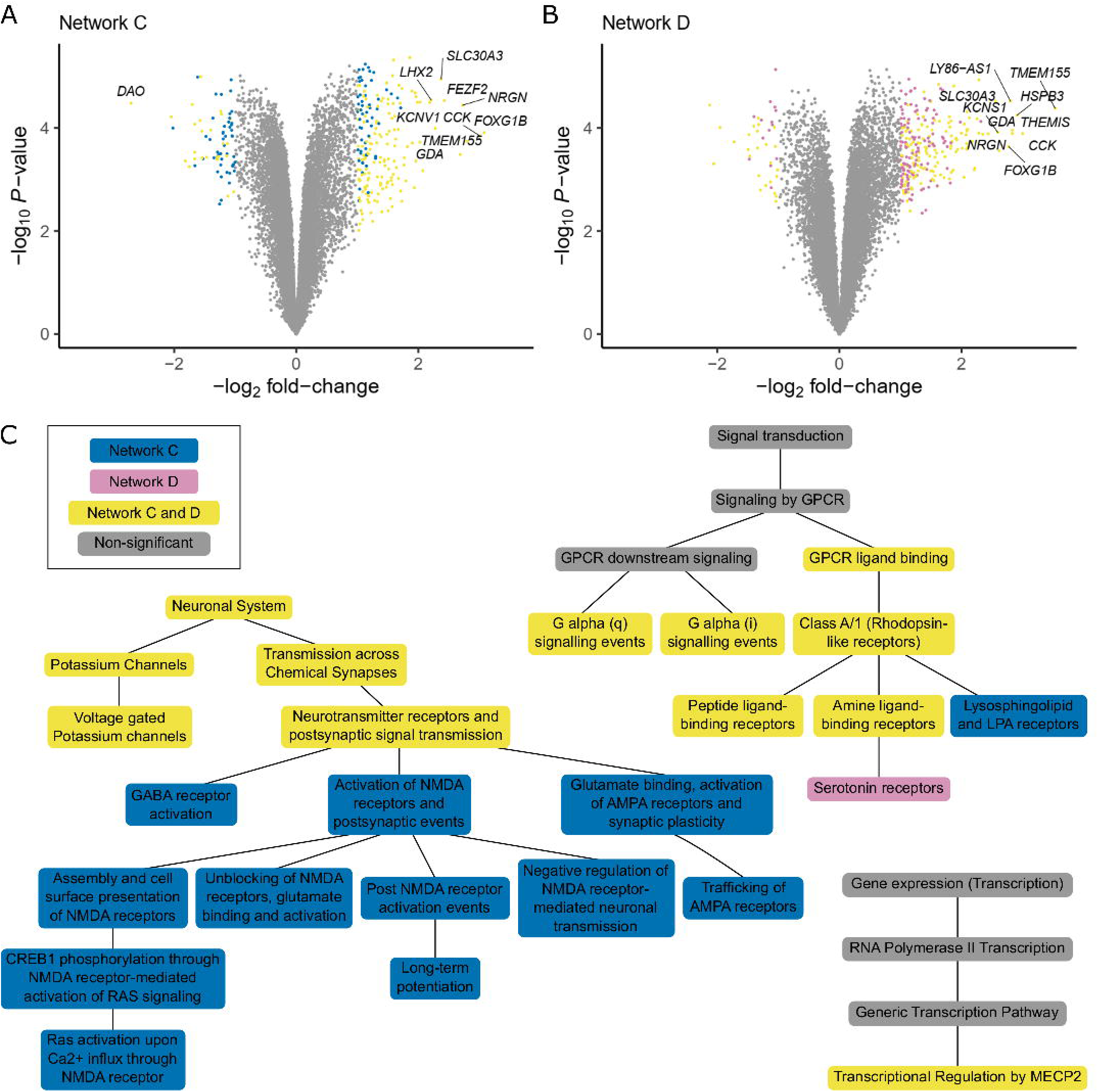
Differential expressed genes and associated pathways. Genes were analyzed for differential expression in the (A) posterior cingulate network (network C) and (B) anterior cingulate network (network D). Effect sizes were summarized across the six healthy donors of AHBA. For all genes (points) the log_2_ fold-change (FC; x-axis) and -log_10_ of nominal *P*-values (y-axis) are shown. Significant differentially expressed genes (t-test, BH-corrected ^P^ < 0.05 and |FC| > 1) are unique for each network (blue and purple points) or significant in both networks (yellow points). For each network the top 10 genes with the highest absolute FC are highlighted, which highly overlap between both networks. (C) Pathway analysis of differentially upregulated genes in the posterior cingulate network and anterior cingulate network. Both networks share similar pathways (yellow) that are hierarchically organized in the Reactome database. The posterior cingulate network showed more specific associations with pathways involved in neurotransmitter receptors and postsynaptic signal transmission (blue). The anterior cingulate network was more specifically associated with serotonin receptors (purple). See Supplementary Table 6 for gene counts and BH-corrected *P*-values.

For functional interpretation, differentially upregulated genes were further assessed for enrichment of genes involved in pathways (Supplementary Table 6). As both networks shared many differentially expressed genes, they also shared similar pathways involved in transcriptional regulation by MECP2, GPCR signaling, voltage gated potassium channels, and neurotransmitter receptor and postsynaptic signal transmission (Figure 2C). The associated pathways were hierarchically related to each other based on the ontology of the Reactome Pathway Database. The posterior cingulate network was additionally related to more specific pathways such as lysosphingolipid and LPA receptors, GABA receptor activation, RAS-signaling mediated by NMDA receptors, glutamate binding, activation of AMPA receptors and synaptic plasticity, and long-term potentiation. The anterior cingulate cortex was additionally associated with serotonin receptors.

### Cell-type enrichment in anatomical networks

The composition of specific cell-types can shape the transcriptomic features of anatomical networks. Therefore, we analyzed whether genes differentially expressed in the posterior and anterior cingulate networks were enriched for cell-type specific marker genes. Based on the average expression of marker genes of a cell-type, the posterior and anterior cingulate networks showed high expression of gene markers for brainstem cholinergic cells, GabaSSTReln, GabaVIPReln, glutamatergic, and pyramidal cells (Figure 3 and Supplementary Figure 1).

**Figure 3.**
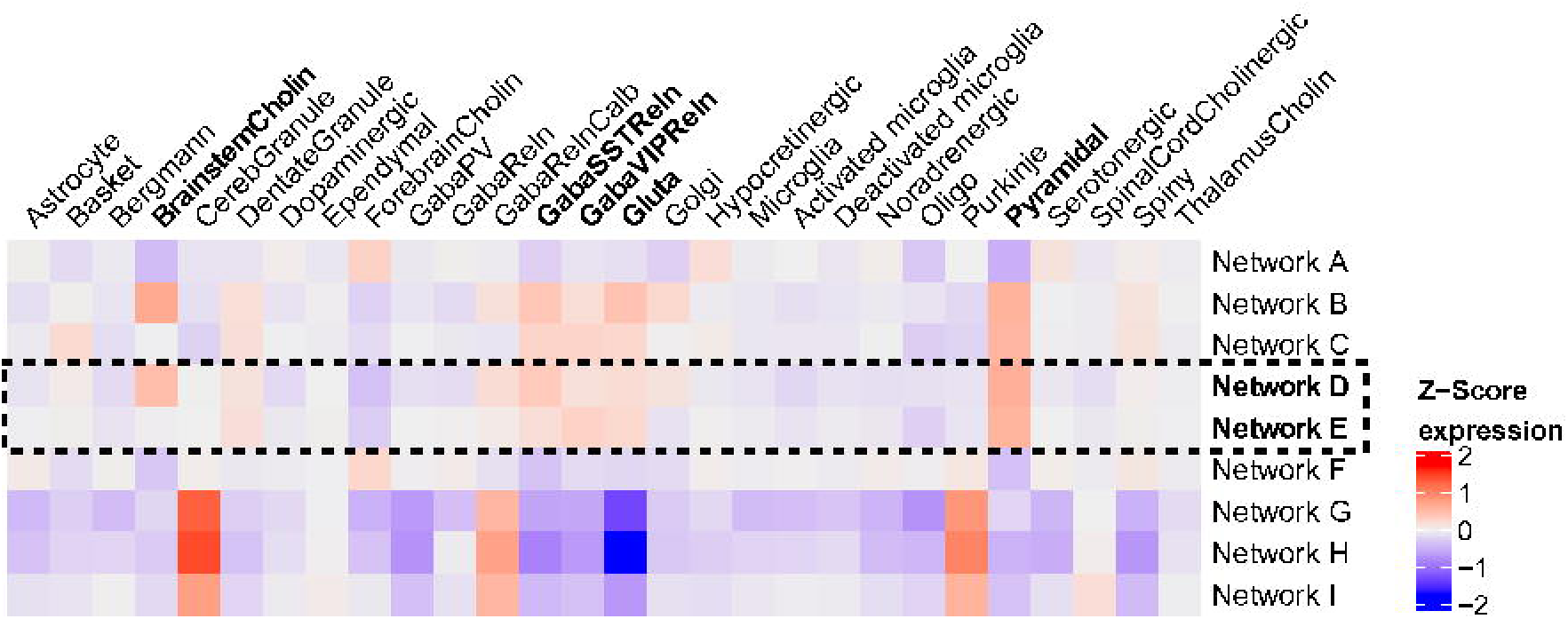
Expression of cell-types in anatomical networks. Gene expression was Z-scored and averaged across cell-type specific markers, samples within anatomical networks, and across the six donors in the Allen Human Brain Atlas. Separate heatmaps for each donor are shown in Supplementary Figure 1.

Among the differentially upregulated genes in the posterior and anterior cingulate networks, we found 10 marker genes representing six cell-types: astrocyte, Bergmann, GabaVIPReln, hypocretinergic, pyramidal, and thalamus cholinergic (Table 2). Those that were significantly upregulated in the posterior cingulate network were also significantly upregulated in the anterior cingulate network. In both networks, the 10 markers were highly expressed in cortical regions, including the cingulate gyrus, and lowly expressed in limbic regions (Figure 4 and Supplementary Figure 2).

**Figure 4.**
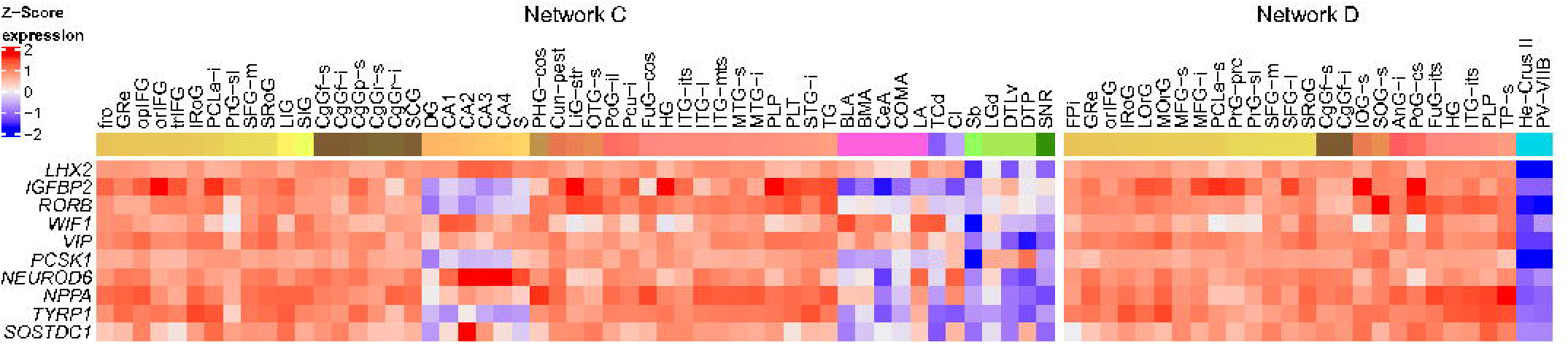
Expression of differentially upregulated cell-type marker genes in the posterior cingulate network (network C) and anterior cingulate network (network D). Heatmaps of differentially expressed marker genes (rows) are shown for one of the six donors in the Allen Human Brain Atlas (donor 10021). Samples from different anatomical substructures within the networks are color annotated (columns). Expression was averaged across samples from an anatomical substructure with the same acronym ignoring left and right hemisphere annotations. See Supplementary Figure 2 for heatmaps for all six donors and Supplementary Table 7 for full names of the region-specific acronyms.

**Table 2.** Differentially upregulated cell-type marker genes in the posterior cingulate network (C) and anterior cingulate network (D). Fold-change (FC) and Benjamini-Hochberg (BH) corrected *P*-value for cell-type markers genes that were differentially expressed in the PD-related networks compared to the non-PD-related networks. FC > 1 and BH < 0.05 are highlighted in red text.

| Gene | Marker | Network C |  | Network D |  |
| --- | --- | --- | --- | --- | --- |
|  |  | FC | BH | Estimate | BH |
| *LHX2* | Astrocyte | 2.21 | 3.92E-03 | 2.00 | 6.46E-03 |
| *IGFBP2* | Astrocyte | 0.69 | 5.80E-02 | 1.18 | 1.78E-02 |
| *RORB* | Astrocyte | 0.82 | 3.09E-02 | 1.19 | 1.39E-02 |
| *WIF1* | Bergmann | 1.02 | 8.74E-03 | 1.03 | 7.95E-03 |
| *VIP* | GabaVIPReIn | 1.67 | 4.23E-03 | 1.85 | 6.89E-03 |
| *PCSK1* | Hypocretinergic | 1.15 | 1.25E-02 | 1.57 | 1.06E-02 |
| *NEUROD6* | Pyramidal | 1.90 | 4.78E-03 | 1.92 | 6.76E-03 |
| *NPPA* | ThalamusCholin | 1.64 | 6.98E-03 | 2.09 | 6.39E-03 |
| *TYRP1* | ThalamusCholin | 0.81 | 2.41E-02 | 1.43 | 9.82E-03 |
| *SOSTDC1* | ThalamusCholin | 0.83 | 1.21E-02 | 1.14 | 6.39E-03 |

Only genes upregulated in the anterior cingulate gyrus were significantly enriched for a cell-type, namely thalamus cholinergic cells (OR = 17.12 and *P* = 2.01e-02). The responsible markers *NPPA, SOSTDC1*, and *TYRP1* showed high expression within the anterior cingulate gyrus network, as well as in most parts of the posterior cingulate gyrus network (Figure 4). Additionally, we showed that their expression was low in limbic samples, including the thalamus, and high in cortical samples within both networks. Interestingly, other thalamus cholinergic marker genes showed opposite expression patterns with high expression in limbic samples and low expression in cortical samples (Supplementary Figure 3).

### PD-related networks are transcriptionally associated to other brain diseases

Dysregulation of functional networks may result in a broader spectrum of disorders than PD. In addition, PD comprises a spectrum of disorders that may result from network dysfunction. Therefore, we assessed which disease-associated genes from DisGeNET were overrepresented among the differentially upregulated genes in the posterior cingulate network as well as the anterior cingulate network. Since both networks shared many upregulated genes, similar disease-associations were also found. We find that genes upregulated in both networks were significantly associated with epileptic and non-epileptic seizures, many mental disorders (bipolar, panic, autistic, cocaine-related, (age-related) memory, mood, major depressive, and anxiety disorder), pain and schizophrenia (Figure 5). The posterior cingulate network was more related to memory and pain-related disorders, while the anterior cingulate network was more related to mental and neuropsychiatric disorders.

**Figure 5.**
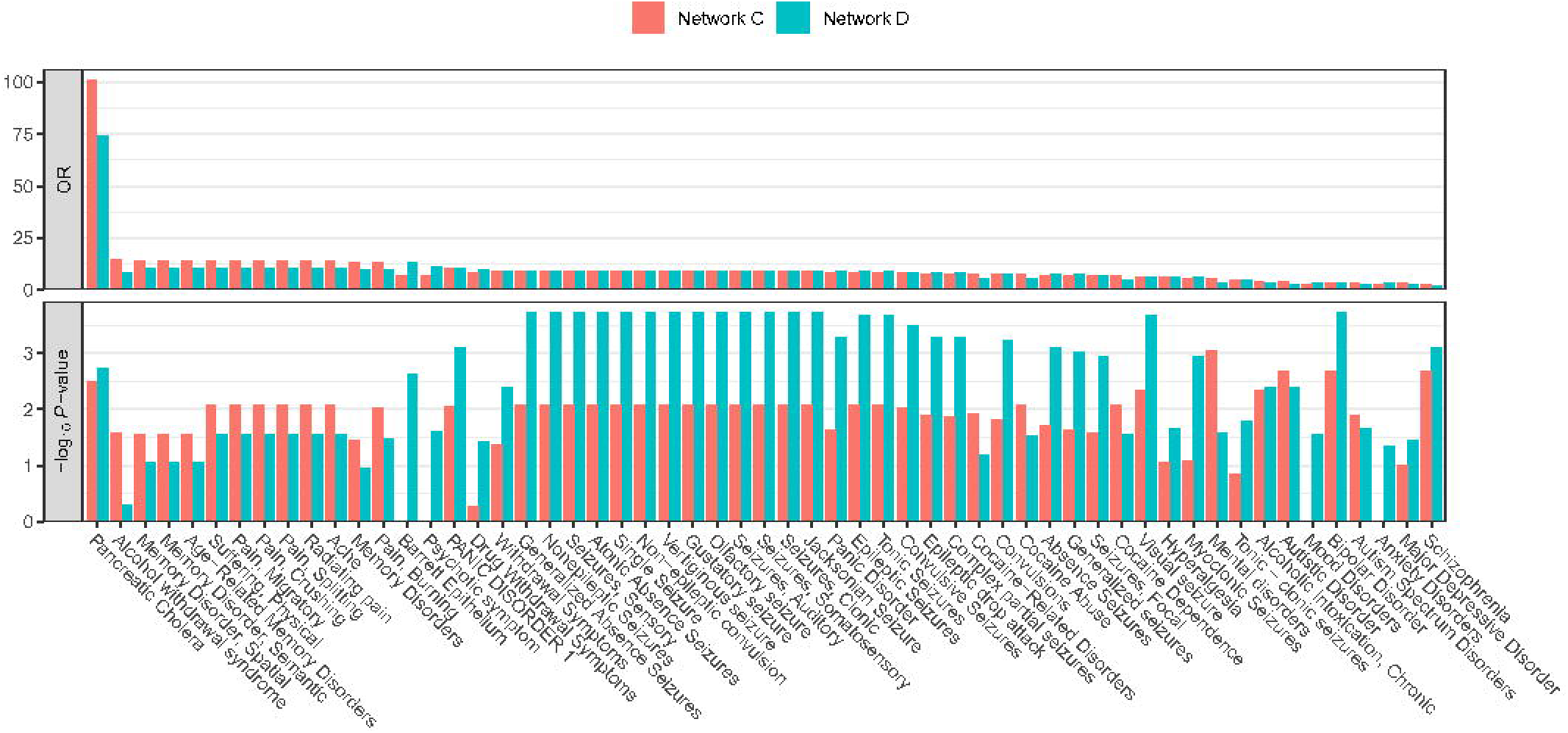
Disease associations of the posterior cingulate network (network C) and anterior cingulate network (network D). Differentially upregulated genes in each network were assessed for the enrichment of disease-associated genes from DisGeNET (hypergeometric test, BH-corrected ^P^ < 0.05). Top plot shows odds ratios (ORs), and bottom plot shows significance of overlap indicated with –log10 *P*-values (y-axis). Disorders (columns) are sorted based on highest ORs in either one of the networks.

## Discussion

The posterior and anterior cingulate networks have been associated earlier with decreased gray matter in PD patients. We examined transcriptomic signatures of both networks in the healthy brain to identify molecular mechanisms underlying gray matter loss in PD. Pathway analysis revealed genes related to gPCR signaling, transcriptional regulation by *MECP2*, and neurotransmitter receptors and postsynaptic signal transmission. We only found significant enrichment of cell-types for genes upregulated in the anterior cingulate gyrus, which were the thalamus cholinergic marker genes. Upon further examination the specific genes were also highly expressed in the posterior cingulate cortex, although not significantly. Moreover, our results showed that SCNs involved in the pathology of PD are associated with multiple neurotransmitter signaling pathways, e.g. serotonin, GPCR, GABA, glutamate, and RAS.

### Cholinergic function in PD

Genes that were highly expressed in the anterior cingulate network were significantly enriched for thalamus cholinergic markers, specifically: *NPPA, SOSTDC1*, and *TYRP1*. These marker genes, together with other markers of this cell-type, were defined based on their expression in cholinergic cells from the thalamus, more specifically the hubenula (Mancarci et al. 2017). Although these genes are considered thalamus-specific, according to the AHBA ontology the hubenula is not part of the thalamus. In this study, most thalamus cholinergic marker genes indeed showed high expression in thalamic regions. However, *NPPA, SOSTDC1*, and *TYRP1*, showed opposite expression patterns with mainly high expression in cortical regions and low expression in limbic regions, including the thalamus. Cholinergic circuits are key in cognitive functions which are impaired in neurodegenerative diseases, such as Alzheimer’s disease and PD (Ballinger et al. 2016). In addition, cholinergic denervation of the cortex and thalamus In PD patients may contribute to the transition from PD to PD with dementia (Ballinger et al. 2016). We also found that glutamatergic and GABAergic marker genes were highly expressed in the AHBA samples within the posterior and anterior cingulate networks, although statistical significance could not be assessed due to the small number of marker genes. Interestingly, acetylcholine release by cholinergic neurons affects glutamatergic and GABAergic signaling by altering the synaptic excitability (Granger et al. 2015; Buendia et al. 2019). Moreover, it is thought that dysfunction of cholinergic circuits contributes to cognitive decline associated with neurodegenerative diseases (Ballinger et al. 2016).

### *NPPA, SOSTDC1*, and TYRP1

The cholinergic marker genes *NPPA, SOSTDC1*, *TYRP1* were highly expressed across the healthy donors in the posterior cingulate network and anterior cingulate network compared to the other seven SCNs. While the functions of these genes likely involve cholinergic signaling, several studies suggest they also function as extracellular regulators of multiple other signaling pathways, including cAMP, Wnt, and β-catenin signaling (Brenner et al. 1990; Hirobe 2011; Kutchko and Siltberg-Liberles 2013; De Vito 2014; Bansho et al. 2017; Millan et al. 2019).

*NPPA* (natriuretic peptide precursor A) and other natriuretic peptides are thought to be involved in a wide range of functions, including neurovascular functions, blood-brain barrier, brain homeostasis, neuroprotection, and synaptic transmission by regulating the release and re-uptake of neurotransmitters such as noradrenalin, dopamine and glycine (Mahinrad et al. 2016). Impaired function of natriuretic peptides in brains of AD patients could accelerate neurodegeneration and may impair structural integrity of the brain leading to a higher risk of cognitive decline (Mahinrad et al. 2018). Our results suggest that *NPPA* might similarly be involved in PD pathogenesis given its high expression within the anterior and posterior cingulate networks.

*SOSTDC1* (sclerostin domain-containing 1) is known as a negative regulator of bone morphogenetic protein (BMP) and Wnt-signaling, but recent studies also show that *SOSTDC1* regulates natural killer cell maturation and cytotoxicity (Millan et al. 2019). An increased number of natural killer cells have been found in PD, but the actual relevance with PD risk is still unclear (Jiang et al. 2017). The BMP signaling pathway promotes the development of midbrain dopaminergic neurons (Jovanovic et al. 2018), in which *SOSTDC1* may play a role. Furthermore, *SOSTDC1* was upregulated in the striatum of Parkinsonian rats that were treated by subthalamic nucleus high frequency stimulation, and is therefore suggested to have neuroprotective effects (Lortet et al. 2013).

*TYRP1* (tyrosinase-related protein 1) produces melanocytes-specific proteins involved in the biosynthesis of melanin in brain, skin and eyes (Wang and Hebert 2006; Lu et al. 2011). Melanoma and PD share genes involved in the synthesis of melanin and dopamine, including *SNCA* which encodes the α-synuclein protein found in Lewy bodies (Pan et al. 2012). Furthermore, neuromelanin is produced almost exclusively in human catecholaminergic neurons and is responsible for the pigmentation of dopaminergic neurons of the substantia nigra, and noradrenergic neurons of the locus cereleus (Pavan and Dalpiaz 2017). It is considered to be protective due to its ability to chelate metals, especially iron which increases with age (Pavan and Dalpiaz 2017).

### Disease-associations

The posterior and anterior cingulate networks shared similar highly expressed genes and were likewise associated with similar diseases. Both SCNs represent anatomical networks that function normally in healthy brains, but their activity is reduced in aging and PD (Hafkemeijer et al. 2014; de Schipper et al. 2017). As part of the default mode network, both the posterior and anterior cingulate cortex have been shown to be dysregulated in neuropsychiatric disorders (Broyd et al. 2009; Öngür et al. 2010). Based on our analysis of transcriptomic signatures in the healthy brain, we found that the posterior cingulate network showed stronger associations with memory and pain-related disorders compared to the anterior cingulate networks which showed stronger associations with mental and neuropsychiatric disorders. Our findings suggest that genes involved in multiple signaling pathways, such as serotonin, GPCR, GABA, glutamate, and RAS, contribute to healthy functions of the posterior and anterior cingulate networks.

## Supplementary material

Supplementary material is available online.

## Acknowledgements

We thank Dr. L. E. Jonkman for her critical insight on the manuscript.

## Notes

*Conflict of Interest:* None declared.

