## Supplementary Materials for "Cholinergic circuit genes in the healthy brain are differentially expressed in regions that exhibit gray matter loss in Parkinson’s disease"

### Supplementary Tables

Supplementary Table 1 Number of cell-type markers. Cell-type markers based on mouse single-cell RNA-sequencing data. After conversion to human homologs and filtering for genes present in AHBA data.

| **Cell-type** | **Mouse** | **Human** | **Cell-type** | **Mouse** | **Human** |
| --- | --- | --- | --- | --- | --- |
| Astrocyte | 97 | 95 | Glutamatergic | 1 | 1 |
| Basket | 1 | 1 | Golgi | 4 | 4 |
| Bergmann | 32 | 30 | Hypocretinergic | 10 | 10 |
| Brainstem cholinergic | 1 | 1 | Microglia | 122 | 120 |
| Cerebellar granule | 9 | 9 | Activated microglia | 88 | 85 |
| Dentate granule | 7 | 7 | Deactivated microglia | 107 | 103 |
| Dopaminergic | 1 | 1 | Noradrenergic | 8 | 8 |
| Ependymal | 45 | 42 | Oligo | 23 | 23 |
| Forebrain cholinergic | 5 | 5 | Purkinje | 27 | 27 |
| GabaPV | 3 | 3 | Pyramidal | 1 | 1 |
| GabaReln | 6 | 6 | Serotonergic | 7 | 7 |
| GabaRelnCalb | 1 | 1 | SpinalCord cholinergic | 7 | 6 |
| GabaSSTReln | 5 | 4 | Spiny | 10 | 10 |
| GabaVIPReln | 5 | 5 | Thalamus cholinergic | 16 | 16 |

Supplementary Table 2 Differentially downregulated genes in the posterior cingulate network (C).

| **Gene** | **Entrez ID** | **Fold-change** | **Lower95** | **Upper95** | ***P*-value** | **BH** |
| --- | --- | --- | --- | --- | --- | --- |
| *ALOX5* | 240 | -1.07 | -1.22 | -0.91 | 1.16E-05 | 3.92E-03 |
| *ANK1* | 286 | -1.06 | -1.26 | -0.85 | 4.76E-05 | 3.92E-03 |
| *CDH7* | 1005 | -1.09 | -1.31 | -0.87 | 5.28E-05 | 3.92E-03 |
| *CHRNA3* | 1136 | -1.08 | -1.31 | -0.85 | 6.69E-05 | 3.92E-03 |
| *DAO* | 1610 | -2.71 | -3.20 | -2.21 | 3.29E-05 | 3.92E-03 |
| *GRM4* | 2914 | -1.09 | -1.32 | -0.86 | 6.40E-05 | 3.92E-03 |
| *KRT19* | 3880 | -1.07 | -1.25 | -0.90 | 1.91E-05 | 3.92E-03 |
| *MEIS1* | 4211 | -1.23 | -1.43 | -1.02 | 2.22E-05 | 3.92E-03 |
| *STON1* | 11037 | -1.23 | -1.47 | -0.98 | 5.37E-05 | 3.92E-03 |
| *RCAN3* | 11123 | -1.32 | -1.54 | -1.09 | 2.19E-05 | 3.92E-03 |
| *RHBG* | 57127 | -1.37 | -1.65 | -1.08 | 6.42E-05 | 3.92E-03 |
| *PCP2* | 126006 | -1.56 | -1.79 | -1.34 | 1.01E-05 | 3.92E-03 |
| *C5orf38* | 153571 | -1.24 | -1.48 | -0.99 | 4.72E-05 | 3.92E-03 |
| *IRX2* | 153572 | -1.70 | -2.02 | -1.39 | 3.50E-05 | 3.92E-03 |
| *C19orf46* | 163183 | -1.58 | -1.91 | -1.24 | 7.13E-05 | 3.92E-03 |
| *ADAMTS18* | 170692 | -1.62 | -1.85 | -1.38 | 1.03E-05 | 3.92E-03 |
| *CRNDE* | 643911 | -2.05 | -2.47 | -1.62 | 6.07E-05 | 3.92E-03 |
| *PGAM2* | 5224 | -1.11 | -1.35 | -0.87 | 7.64E-05 | 3.98E-03 |
| *C7orf16* | 10842 | -1.06 | -1.29 | -0.82 | 8.05E-05 | 3.99E-03 |
| *IRX5* | 10265 | -1.66 | -2.03 | -1.29 | 8.44E-05 | 4.05E-03 |
| *LDLRAP1* | 26119 | -1.06 | -1.30 | -0.82 | 8.63E-05 | 4.08E-03 |
| *IL28RA* | 163702 | -1.18 | -1.44 | -0.91 | 9.28E-05 | 4.14E-03 |
| *CBLN1* | 869 | -2.02 | -2.48 | -1.55 | 1.01E-04 | 4.19E-03 |
| *IL16* | 3603 | -1.09 | -1.34 | -0.84 | 9.66E-05 | 4.19E-03 |
| *GPRIN2* | 9721 | -1.40 | -1.73 | -1.07 | 1.09E-04 | 4.24E-03 |
| *CALB2* | 794 | -1.06 | -1.31 | -0.81 | 1.11E-04 | 4.25E-03 |
| *EPB41* | 2035 | -1.20 | -1.48 | -0.91 | 1.21E-04 | 4.25E-03 |
| *IRX3* | 79191 | -1.55 | -1.91 | -1.18 | 1.15E-04 | 4.25E-03 |
| *SPINK6* | 404203 | -1.20 | -1.49 | -0.91 | 1.26E-04 | 4.34E-03 |
| *ZNF521* | 25925 | -1.05 | -1.31 | -0.79 | 1.43E-04 | 4.50E-03 |
| *KLHL1* | 57626 | -1.04 | -1.30 | -0.79 | 1.44E-04 | 4.50E-03 |
| *DOK7* | 285489 | -1.48 | -1.85 | -1.11 | 1.46E-04 | 4.50E-03 |
| *CNPY1* | 285888 | -1.16 | -1.45 | -0.87 | 1.45E-04 | 4.50E-03 |
| *THRSP* | 7069 | -1.20 | -1.50 | -0.89 | 1.63E-04 | 4.62E-03 |
| *ZIC3* | 7547 | -1.41 | -1.78 | -1.05 | 1.68E-04 | 4.63E-03 |
| *STAC* | 6769 | -1.20 | -1.52 | -0.88 | 2.02E-04 | 4.82E-03 |
| *TIMP4* | 7079 | -1.13 | -1.43 | -0.83 | 2.02E-04 | 4.82E-03 |
| *PDZK1* | 5174 | -1.18 | -1.50 | -0.86 | 2.18E-04 | 4.95E-03 |
| *LGR6* | 59352 | -1.29 | -1.66 | -0.92 | 2.94E-04 | 5.43E-03 |
| *TFAP2B* | 7021 | -1.66 | -2.14 | -1.18 | 2.94E-04 | 5.43E-03 |
| *CBLN3* | 643866 | -1.24 | -1.60 | -0.88 | 3.15E-04 | 5.55E-03 |
| *LEMD1* | 93273 | -1.14 | -1.47 | -0.80 | 3.19E-04 | 5.58E-03 |
| *EOMES* | 8320 | -1.27 | -1.65 | -0.90 | 3.28E-04 | 5.59E-03 |
| *PIRT* | 644139 | -1.14 | -1.48 | -0.80 | 3.48E-04 | 5.75E-03 |
| *MYT1* | 4661 | -1.02 | -1.33 | -0.72 | 3.59E-04 | 5.80E-03 |
| *SLC35F4* | 341880 | -1.27 | -1.65 | -0.89 | 3.60E-04 | 5.80E-03 |
| *CCDC155* | 147872 | -1.29 | -1.68 | -0.90 | 3.64E-04 | 5.83E-03 |
| *C9orf171* | 389799 | -1.29 | -1.68 | -0.90 | 3.67E-04 | 5.83E-03 |
| *BARHL1* | 56751 | -1.47 | -1.91 | -1.02 | 3.70E-04 | 5.86E-03 |
| *UNCX* | 340260 | -1.35 | -1.76 | -0.94 | 3.80E-04 | 5.92E-03 |
| *SLITRK6* | 84189 | -1.54 | -2.01 | -1.07 | 3.84E-04 | 5.92E-03 |
| *GABRA6* | 2559 | -1.53 | -1.99 | -1.06 | 3.94E-04 | 5.99E-03 |
| *ZIC4* | 84107 | -1.66 | -2.17 | -1.15 | 3.97E-04 | 6.00E-03 |
| *EBF1* | 1879 | -1.22 | -1.60 | -0.85 | 4.02E-04 | 6.03E-03 |
| *MAB21L1* | 4081 | -1.30 | -1.70 | -0.90 | 4.07E-04 | 6.03E-03 |
| *CDH15* | 1013 | -1.82 | -2.39 | -1.25 | 4.36E-04 | 6.20E-03 |
| *NUAK2* | 81788 | -1.00 | -1.32 | -0.68 | 4.64E-04 | 6.33E-03 |
| *IRF6* | 3664 | -1.03 | -1.37 | -0.70 | 5.15E-04 | 6.62E-03 |
| *ANGPTL7* | 10218 | -1.03 | -1.37 | -0.70 | 5.20E-04 | 6.66E-03 |
| *BARHL2* | 343472 | -1.24 | -1.64 | -0.83 | 5.22E-04 | 6.67E-03 |
| *CRTAM* | 56253 | -1.66 | -2.20 | -1.12 | 5.39E-04 | 6.72E-03 |
| *SYT2* | 127833 | -1.04 | -1.39 | -0.70 | 5.48E-04 | 6.77E-03 |
| *ZIC1* | 7545 | -1.75 | -2.33 | -1.17 | 5.61E-04 | 6.82E-03 |
| *STK32A* | 202374 | -1.02 | -1.36 | -0.67 | 6.81E-04 | 7.48E-03 |
| *PCSK9* | 255738 | -1.29 | -1.74 | -0.84 | 7.12E-04 | 7.64E-03 |
| *ZP2* | 7783 | -1.14 | -1.53 | -0.74 | 7.29E-04 | 7.70E-03 |
| *SLC22A31* | 146429 | -1.09 | -1.47 | -0.71 | 7.57E-04 | 7.82E-03 |
| *CLCNKB* | 1188 | -1.23 | -1.68 | -0.78 | 8.91E-04 | 8.40E-03 |
| *FLT3* | 2322 | -1.10 | -1.52 | -0.67 | 1.17E-03 | 9.56E-03 |
| *DLK1* | 8788 | -1.03 | -1.46 | -0.59 | 1.73E-03 | 1.18E-02 |
| *EBF3* | 253738 | -1.12 | -1.62 | -0.63 | 2.10E-03 | 1.31E-02 |
| *FAT2* | 2196 | -1.23 | -1.79 | -0.66 | 2.52E-03 | 1.45E-02 |
| *PVALB* | 5816 | -1.26 | -1.86 | -0.65 | 3.01E-03 | 1.62E-02 |

Supplementary Table 3 Differentially downregulated genes in the anterior cingulate network (D).

| **Gene** | **Entrez ID** | **Fold-change** | **Lower95** | **Upper95** | ***P*-value** | **BH** |
| --- | --- | --- | --- | --- | --- | --- |
| *CA12* | 771 | -1.40 | -1.69 | -1.11 | 6.02E-05 | 6.39E-03 |
| *CHRNA3* | 1136 | -1.33 | -1.60 | -1.05 | 6.41E-05 | 6.39E-03 |
| *DRD2* | 1813 | -1.05 | -1.19 | -0.90 | 7.31E-06 | 6.39E-03 |
| *MAB21L1* | 4081 | -1.06 | -1.31 | -0.82 | 9.63E-05 | 6.39E-03 |
| *PENK* | 5179 | -1.11 | -1.35 | -0.86 | 7.67E-05 | 6.39E-03 |
| *STAC* | 6769 | -1.32 | -1.61 | -1.03 | 8.59E-05 | 6.39E-03 |
| *LIPG* | 9388 | -1.15 | -1.42 | -0.88 | 1.11E-04 | 6.39E-03 |
| *GPRIN2* | 9721 | -1.34 | -1.66 | -1.03 | 1.15E-04 | 6.39E-03 |
| *RGS8* | 85397 | -1.28 | -1.55 | -1.00 | 7.00E-05 | 6.39E-03 |
| *IRX2* | 153572 | -1.96 | -2.40 | -1.51 | 9.72E-05 | 6.39E-03 |
| *C19orf46* | 163183 | -1.40 | -1.72 | -1.09 | 9.22E-05 | 6.39E-03 |
| *DACT2* | 168002 | -1.06 | -1.26 | -0.86 | 3.93E-05 | 6.39E-03 |
| *ANKRD34B* | 340120 | -1.10 | -1.34 | -0.85 | 8.65E-05 | 6.39E-03 |
| *RSPO4* | 343637 | -1.04 | -1.25 | -0.84 | 4.59E-05 | 6.39E-03 |
| *CRNDE* | 643911 | -2.12 | -2.52 | -1.73 | 3.60E-05 | 6.39E-03 |
| *TMEM90A* | 646658 | -1.13 | -1.33 | -0.93 | 3.08E-05 | 6.39E-03 |
| *IRX3* | 79191 | -1.48 | -1.84 | -1.12 | 1.35E-04 | 6.54E-03 |
| *ECEL1* | 9427 | -1.25 | -1.57 | -0.94 | 1.57E-04 | 6.76E-03 |
| *IRX5* | 10265 | -1.72 | -2.18 | -1.27 | 1.87E-04 | 6.90E-03 |
| *ZIC4* | 84107 | -1.20 | -1.52 | -0.88 | 2.09E-04 | 6.93E-03 |
| *SLC35F4* | 341880 | -1.05 | -1.34 | -0.76 | 2.28E-04 | 7.13E-03 |
| *SLITRK6* | 84189 | -1.34 | -1.71 | -0.97 | 2.44E-04 | 7.22E-03 |
| *C5orf38* | 153571 | -1.49 | -1.91 | -1.07 | 2.62E-04 | 7.35E-03 |
| *CDH15* | 1013 | -1.47 | -1.91 | -1.03 | 3.54E-04 | 7.95E-03 |
| *COL8A2* | 1296 | -1.03 | -1.34 | -0.71 | 3.95E-04 | 8.20E-03 |
| *NTS* | 4922 | -1.47 | -1.93 | -1.02 | 4.03E-04 | 8.23E-03 |
| *DLK1* | 8788 | -1.13 | -1.48 | -0.77 | 4.77E-04 | 8.75E-03 |
| *DAO* | 1610 | -2.06 | -2.72 | -1.40 | 4.94E-04 | 8.87E-03 |
| *WNT11* | 7481 | -1.13 | -1.49 | -0.76 | 5.39E-04 | 9.08E-03 |
| *PIRT* | 644139 | -1.01 | -1.36 | -0.67 | 6.54E-04 | 9.91E-03 |
| *C9orf171* | 389799 | -1.32 | -1.79 | -0.84 | 8.43E-04 | 1.06E-02 |
| *TIMP4* | 7079 | -1.02 | -1.40 | -0.65 | 8.71E-04 | 1.07E-02 |
| *STON1* | 11037 | -1.13 | -1.56 | -0.70 | 1.11E-03 | 1.20E-02 |
| *TFAP2B* | 7021 | -1.20 | -1.68 | -0.73 | 1.32E-03 | 1.30E-02 |
| *GBX2* | 2637 | -1.12 | -1.57 | -0.66 | 1.45E-03 | 1.36E-02 |
| *PCP2* | 126006 | -1.29 | -1.83 | -0.76 | 1.61E-03 | 1.42E-02 |
| *EBF3* | 253738 | -1.06 | -1.51 | -0.60 | 1.86E-03 | 1.53E-02 |
| *ZIC1* | 7545 | -1.04 | -1.50 | -0.58 | 2.12E-03 | 1.63E-02 |
| *BARHL1* | 56751 | -1.04 | -1.52 | -0.56 | 2.65E-03 | 1.83E-02 |

Supplementary Table 4 Differentially upregulated genes in the posterior cingulate network (C).

| **Gene** | **Entrez ID** | **Fold-change** | **Lower95** | **Upper95** | ***P*-value** | **BH** |
| --- | --- | --- | --- | --- | --- | --- |
| *ADRA1D* | 146 | 1.33 | 1.07 | 1.58 | 4.27E-05 | 3.92E-03 |
| *CAMK2A* | 815 | 1.95 | 1.62 | 2.27 | 2.08E-05 | 3.92E-03 |
| *CDH9* | 1007 | 1.68 | 1.42 | 1.94 | 1.48E-05 | 3.92E-03 |
| *CHN1* | 1123 | 1.13 | 0.90 | 1.36 | 5.88E-05 | 3.92E-03 |
| *EGR3* | 1960 | 2.03 | 1.66 | 2.39 | 3.14E-05 | 3.92E-03 |
| *EXTL1* | 2134 | 1.16 | 0.94 | 1.38 | 3.90E-05 | 3.92E-03 |
| *F12* | 2161 | 1.05 | 0.87 | 1.22 | 2.27E-05 | 3.92E-03 |
| *FHL2* | 2274 | 1.50 | 1.24 | 1.77 | 2.75E-05 | 3.92E-03 |
| *GABRA5* | 2558 | 1.98 | 1.62 | 2.34 | 3.17E-05 | 3.92E-03 |
| *GRIN2B* | 2904 | 1.21 | 0.98 | 1.44 | 4.07E-05 | 3.92E-03 |
| *HTR1A* | 3350 | 1.33 | 1.10 | 1.55 | 2.31E-05 | 3.92E-03 |
| *ITPKA* | 3706 | 1.34 | 1.09 | 1.59 | 3.78E-05 | 3.92E-03 |
| *LMO7* | 4008 | 1.00 | 0.80 | 1.20 | 4.84E-05 | 3.92E-03 |
| *NELL2* | 4753 | 1.01 | 0.83 | 1.18 | 2.48E-05 | 3.92E-03 |
| *NNMT* | 4837 | 1.39 | 1.14 | 1.64 | 3.00E-05 | 3.92E-03 |
| *NPTX2* | 4885 | 1.50 | 1.23 | 1.78 | 3.41E-05 | 3.92E-03 |
| *NRGN* | 4900 | 2.73 | 2.22 | 3.23 | 3.61E-05 | 3.92E-03 |
| *PCDH8* | 5100 | 1.70 | 1.39 | 2.01 | 3.30E-05 | 3.92E-03 |
| *PDE2A* | 5138 | 1.59 | 1.40 | 1.79 | 4.78E-06 | 3.92E-03 |
| *PSD* | 5662 | 1.02 | 0.88 | 1.17 | 1.07E-05 | 3.92E-03 |
| *RAB27B* | 5874 | 1.15 | 0.92 | 1.38 | 4.97E-05 | 3.92E-03 |
| *RASGRF2* | 5924 | 1.04 | 0.88 | 1.20 | 1.49E-05 | 3.92E-03 |
| *SLC30A3* | 7781 | 2.37 | 2.02 | 2.72 | 1.12E-05 | 3.92E-03 |
| *ENC1* | 8507 | 1.96 | 1.62 | 2.30 | 2.46E-05 | 3.92E-03 |
| *LMO4* | 8543 | 1.02 | 0.80 | 1.24 | 6.85E-05 | 3.92E-03 |
| *HRK* | 8739 | 1.27 | 1.09 | 1.44 | 8.33E-06 | 3.92E-03 |
| *KALRN* | 8997 | 1.13 | 0.98 | 1.27 | 5.77E-06 | 3.92E-03 |
| *LDB2* | 9079 | 1.73 | 1.45 | 2.01 | 1.71E-05 | 3.92E-03 |
| *LHX2* | 9355 | 2.21 | 1.82 | 2.60 | 2.86E-05 | 3.92E-03 |
| *CARTPT* | 9607 | 1.71 | 1.38 | 2.04 | 4.08E-05 | 3.92E-03 |
| *CACNG3* | 10368 | 1.52 | 1.22 | 1.83 | 5.19E-05 | 3.92E-03 |
| *PLK2* | 10769 | 1.48 | 1.24 | 1.72 | 1.81E-05 | 3.92E-03 |
| *LZTS1* | 11178 | 1.48 | 1.26 | 1.71 | 1.33E-05 | 3.92E-03 |
| *MAST3* | 23031 | 1.20 | 1.04 | 1.37 | 7.91E-06 | 3.92E-03 |
| *KCNH3* | 23416 | 1.11 | 0.93 | 1.30 | 1.93E-05 | 3.92E-03 |
| *RIMBP2* | 23504 | 1.03 | 0.89 | 1.17 | 7.36E-06 | 3.92E-03 |
| *MMD* | 23531 | 1.13 | 0.94 | 1.32 | 2.34E-05 | 3.92E-03 |
| *AK5* | 26289 | 1.61 | 1.28 | 1.95 | 5.90E-05 | 3.92E-03 |
| *SYT17* | 51760 | 1.08 | 0.86 | 1.30 | 5.34E-05 | 3.92E-03 |
| *SEMA5B* | 54437 | 1.04 | 0.88 | 1.19 | 1.24E-05 | 3.92E-03 |
| *FEZF2* | 55079 | 2.43 | 1.99 | 2.86 | 2.94E-05 | 3.92E-03 |
| *KCNQ5* | 56479 | 1.14 | 0.94 | 1.34 | 2.72E-05 | 3.92E-03 |
| *SLC17A7* | 57030 | 1.90 | 1.58 | 2.21 | 1.93E-05 | 3.92E-03 |
| *CAMK1G* | 57172 | 1.03 | 0.83 | 1.23 | 4.50E-05 | 3.92E-03 |
| *KIAA1324* | 57535 | 1.07 | 0.91 | 1.23 | 1.19E-05 | 3.92E-03 |
| *LRRC7* | 57554 | 1.66 | 1.37 | 1.95 | 2.62E-05 | 3.92E-03 |
| *CLSTN2* | 64084 | 1.02 | 0.83 | 1.21 | 3.43E-05 | 3.92E-03 |
| *PCDH20* | 64881 | 1.57 | 1.24 | 1.89 | 6.48E-05 | 3.92E-03 |
| *AC006273.1* | 79948 | 1.23 | 1.05 | 1.41 | 1.19E-05 | 3.92E-03 |
| *NETO1* | 81832 | 1.59 | 1.30 | 1.88 | 3.36E-05 | 3.92E-03 |
| *ST6GALNAC5* | 81849 | 1.58 | 1.35 | 1.80 | 9.67E-06 | 3.92E-03 |
| *SYT16* | 83851 | 1.08 | 0.88 | 1.27 | 3.05E-05 | 3.92E-03 |
| *NCALD* | 83988 | 1.10 | 0.91 | 1.29 | 2.40E-05 | 3.92E-03 |
| *SYDE2* | 84144 | 1.04 | 0.82 | 1.26 | 6.81E-05 | 3.92E-03 |
| *LRRC62* | 114794 | 1.05 | 0.90 | 1.21 | 1.03E-05 | 3.92E-03 |
| *CPNE4* | 131034 | 1.56 | 1.30 | 1.82 | 2.18E-05 | 3.92E-03 |
| *LOC158696* | 158696 | 1.13 | 0.96 | 1.31 | 1.51E-05 | 3.92E-03 |
| *KCNG3* | 170850 | 1.34 | 1.11 | 1.58 | 2.75E-05 | 3.92E-03 |
| *CREG2* | 200407 | 2.18 | 1.79 | 2.58 | 3.18E-05 | 3.92E-03 |
| *C8orf46* | 254778 | 1.09 | 0.95 | 1.24 | 7.27E-06 | 3.92E-03 |
| *CHSY3* | 337876 | 1.18 | 1.03 | 1.34 | 6.67E-06 | 3.92E-03 |
| *FAM19A2* | 338811 | 1.20 | 0.97 | 1.42 | 3.69E-05 | 3.92E-03 |
| *MYBPHL* | 343263 | 1.20 | 0.98 | 1.42 | 3.54E-05 | 3.92E-03 |
| *KCNT2* | 343450 | 1.01 | 0.85 | 1.17 | 1.50E-05 | 3.92E-03 |
| *C2orf55* | 343990 | 1.42 | 1.17 | 1.67 | 2.65E-05 | 3.92E-03 |
| *C1orf95* | 375057 | 1.08 | 0.88 | 1.28 | 3.67E-05 | 3.92E-03 |
| *KCTD4* | 386618 | 1.24 | 0.98 | 1.50 | 6.31E-05 | 3.92E-03 |
| *FAM19A1* | 407738 | 1.86 | 1.63 | 2.08 | 4.29E-06 | 3.92E-03 |
| *LOC440084* | 440084 | 1.06 | 0.85 | 1.26 | 4.47E-05 | 3.92E-03 |
| *CHRM3* | 1131 | 1.32 | 1.03 | 1.60 | 7.38E-05 | 3.94E-03 |
| *KIRREL2* | 84063 | 1.16 | 0.91 | 1.41 | 7.42E-05 | 3.94E-03 |
| *STX1A* | 6804 | 1.11 | 0.87 | 1.35 | 7.69E-05 | 3.98E-03 |
| *DLGAP2* | 9228 | 1.32 | 1.03 | 1.61 | 8.04E-05 | 3.99E-03 |
| *AKAP5* | 9495 | 1.34 | 1.05 | 1.64 | 7.93E-05 | 3.99E-03 |
| *TNFAIP8L3* | 388121 | 1.35 | 1.05 | 1.65 | 8.21E-05 | 4.01E-03 |
| *NEURL1B* | 54492 | 1.23 | 0.95 | 1.51 | 9.11E-05 | 4.14E-03 |
| *RP4-788L13.1* | 9890 | 1.04 | 0.80 | 1.28 | 9.80E-05 | 4.19E-03 |
| *KCNV1* | 27012 | 2.28 | 1.75 | 2.80 | 1.02E-04 | 4.19E-03 |
| *MKL2* | 57496 | 1.07 | 0.82 | 1.32 | 1.01E-04 | 4.19E-03 |
| *MUM1L1* | 139221 | 1.30 | 1.00 | 1.61 | 1.02E-04 | 4.19E-03 |
| *VIP* | 7432 | 1.67 | 1.28 | 2.06 | 1.06E-04 | 4.23E-03 |
| *LY6H* | 4062 | 1.61 | 1.23 | 1.98 | 1.08E-04 | 4.24E-03 |
| *CAMKV* | 79012 | 1.52 | 1.16 | 1.88 | 1.15E-04 | 4.25E-03 |
| *FOXG1B* | 2290 | 3.08 | 2.34 | 3.82 | 1.23E-04 | 4.28E-03 |
| *MPPED1* | 758 | 1.29 | 0.98 | 1.60 | 1.29E-04 | 4.37E-03 |
| *TBR1* | 10716 | 1.49 | 1.12 | 1.85 | 1.32E-04 | 4.38E-03 |
| *DACH2* | 117154 | 1.37 | 1.03 | 1.70 | 1.33E-04 | 4.41E-03 |
| *CCK* | 885 | 3.01 | 2.26 | 3.75 | 1.42E-04 | 4.50E-03 |
| *SCN3B* | 55800 | 1.19 | 0.89 | 1.48 | 1.43E-04 | 4.50E-03 |
| *KCTD16* | 57528 | 1.39 | 1.04 | 1.74 | 1.48E-04 | 4.50E-03 |
| *BAIAP3* | 8938 | 1.20 | 0.90 | 1.50 | 1.52E-04 | 4.52E-03 |
| *ANO3* | 63982 | 1.63 | 1.21 | 2.04 | 1.64E-04 | 4.63E-03 |
| *PPAPR5* | 163404 | 1.07 | 0.80 | 1.35 | 1.71E-04 | 4.65E-03 |
| *TMEM155* | 132332 | 2.80 | 2.07 | 3.52 | 1.79E-04 | 4.74E-03 |
| *RASAL1* | 8437 | 1.15 | 0.85 | 1.45 | 1.84E-04 | 4.77E-03 |
| *GABRA4* | 2557 | 1.13 | 0.83 | 1.43 | 1.94E-04 | 4.78E-03 |
| *ARC* | 23237 | 1.32 | 0.97 | 1.67 | 1.94E-04 | 4.78E-03 |
| *MOXD1* | 26002 | 2.02 | 1.49 | 2.56 | 1.89E-04 | 4.78E-03 |
| *NEUROD6* | 63974 | 1.90 | 1.40 | 2.40 | 1.95E-04 | 4.78E-03 |
| *KHDRBS2* | 202559 | 1.22 | 0.89 | 1.54 | 1.98E-04 | 4.81E-03 |
| *GPR26* | 2849 | 1.49 | 1.09 | 1.89 | 2.05E-04 | 4.87E-03 |
| *C2orf80* | 389073 | 1.23 | 0.90 | 1.56 | 2.06E-04 | 4.87E-03 |
| *HGF* | 3082 | 1.08 | 0.79 | 1.38 | 2.26E-04 | 4.97E-03 |
| *KIAA1239* | 57495 | 1.02 | 0.74 | 1.29 | 2.25E-04 | 4.97E-03 |
| *C14orf23* | 387978 | 1.61 | 1.18 | 2.05 | 2.22E-04 | 4.97E-03 |
| *ICAM5* | 7087 | 1.82 | 1.32 | 2.32 | 2.31E-04 | 4.98E-03 |
| *AP003108.2* | 390205 | 1.14 | 0.82 | 1.46 | 2.52E-04 | 5.16E-03 |
| *C1orf115* | 79762 | 1.21 | 0.87 | 1.55 | 2.61E-04 | 5.23E-03 |
| *EFNB3* | 1949 | 1.01 | 0.73 | 1.30 | 2.65E-04 | 5.25E-03 |
| *LINGO1* | 84894 | 1.06 | 0.76 | 1.36 | 2.85E-04 | 5.39E-03 |
| *MEF2C* | 4208 | 1.16 | 0.82 | 1.50 | 3.09E-04 | 5.50E-03 |
| *TMEM132B* | 114795 | 1.06 | 0.75 | 1.36 | 3.10E-04 | 5.51E-03 |
| *GDA* | 9615 | 2.68 | 1.89 | 3.47 | 3.26E-04 | 5.59E-03 |
| *ATRNL1* | 26033 | 1.18 | 0.84 | 1.53 | 3.20E-04 | 5.59E-03 |
| *OTX1* | 5013 | 1.14 | 0.80 | 1.48 | 3.37E-04 | 5.63E-03 |
| *THRB* | 7068 | 1.16 | 0.81 | 1.50 | 3.38E-04 | 5.63E-03 |
| *C6orf126* | 389383 | 1.10 | 0.78 | 1.43 | 3.38E-04 | 5.63E-03 |
| *FILIP1* | 27145 | 1.30 | 0.91 | 1.69 | 3.56E-04 | 5.79E-03 |
| *NR2E1* | 7101 | 1.38 | 0.96 | 1.79 | 3.73E-04 | 5.88E-03 |
| *NPY* | 4852 | 1.77 | 1.23 | 2.31 | 3.79E-04 | 5.92E-03 |
| *KLK7* | 5650 | 1.13 | 0.79 | 1.47 | 3.81E-04 | 5.92E-03 |
| *RGS14* | 10636 | 1.30 | 0.90 | 1.70 | 3.89E-04 | 5.95E-03 |
| *SSTR1* | 6751 | 1.02 | 0.71 | 1.33 | 3.96E-04 | 6.00E-03 |
| *FAM81A* | 145773 | 1.51 | 1.05 | 1.97 | 3.99E-04 | 6.01E-03 |
| *KCNS2* | 3788 | 1.11 | 0.77 | 1.45 | 4.05E-04 | 6.03E-03 |
| *NEK2* | 4751 | 1.08 | 0.75 | 1.41 | 4.10E-04 | 6.04E-03 |
| *RGS4* | 5999 | 1.44 | 1.00 | 1.89 | 4.15E-04 | 6.06E-03 |
| *NPTXR* | 23467 | 1.03 | 0.71 | 1.35 | 4.33E-04 | 6.18E-03 |
| *ZNF831* | 128611 | 1.60 | 1.10 | 2.11 | 4.41E-04 | 6.21E-03 |
| *OR14I1* | 401994 | 1.96 | 1.35 | 2.57 | 4.41E-04 | 6.21E-03 |
| *C1QL1* | 389941 | 1.44 | 0.99 | 1.89 | 4.48E-04 | 6.25E-03 |
| *EMX2OS* | 196047 | 1.37 | 0.94 | 1.80 | 4.62E-04 | 6.33E-03 |
| *HRH1* | 3269 | 1.27 | 0.87 | 1.68 | 4.65E-04 | 6.33E-03 |
| *WDR86* | 349136 | 1.08 | 0.74 | 1.43 | 4.74E-04 | 6.38E-03 |
| *EFCAB1* | 79645 | 1.09 | 0.74 | 1.44 | 4.79E-04 | 6.39E-03 |
| *SST* | 6750 | 1.76 | 1.19 | 2.34 | 5.25E-04 | 6.67E-03 |
| *KCNC2* | 3747 | 1.45 | 0.98 | 1.93 | 5.39E-04 | 6.72E-03 |
| *EGR2* | 1959 | 1.15 | 0.77 | 1.53 | 5.45E-04 | 6.76E-03 |
| *NPPA* | 4878 | 1.64 | 1.09 | 2.19 | 5.88E-04 | 6.98E-03 |
| *RBP4* | 5950 | 1.52 | 1.00 | 2.03 | 6.28E-04 | 7.17E-03 |
| *STYK1* | 55359 | 1.15 | 0.76 | 1.53 | 6.32E-04 | 7.17E-03 |
| *FAM148C* | 126567 | 1.01 | 0.67 | 1.35 | 6.37E-04 | 7.20E-03 |
| *KIF17* | 57576 | 1.05 | 0.69 | 1.41 | 6.53E-04 | 7.32E-03 |
| *HSPB3* | 8988 | 2.07 | 1.36 | 2.78 | 6.71E-04 | 7.42E-03 |
| *CDH8* | 1006 | 1.08 | 0.70 | 1.46 | 7.38E-04 | 7.74E-03 |
| *DLX6-AS1* | 285987 | 1.19 | 0.77 | 1.61 | 7.49E-04 | 7.79E-03 |
| *SERPINF1* | 5176 | 1.12 | 0.72 | 1.52 | 7.78E-04 | 7.88E-03 |
| *SLIT1* | 6585 | 1.59 | 1.02 | 2.15 | 7.97E-04 | 8.00E-03 |
| *CORT* | 1325 | 1.08 | 0.69 | 1.46 | 8.12E-04 | 8.05E-03 |
| *TNNT2* | 7139 | 1.33 | 0.84 | 1.81 | 8.95E-04 | 8.40E-03 |
| *DDN* | 23109 | 1.40 | 0.89 | 1.91 | 8.96E-04 | 8.40E-03 |
| *TAC3* | 6866 | 1.56 | 0.99 | 2.13 | 9.16E-04 | 8.48E-03 |
| *LMO3* | 55885 | 1.19 | 0.75 | 1.62 | 9.40E-04 | 8.58E-03 |
| *AC078937.4* | 286002 | 1.67 | 1.05 | 2.29 | 9.60E-04 | 8.67E-03 |
| *WIF1* | 11197 | 1.02 | 0.64 | 1.41 | 9.78E-04 | 8.74E-03 |
| *CCDC3* | 83643 | 1.04 | 0.65 | 1.43 | 1.02E-03 | 8.96E-03 |
| *FAM5B* | 57795 | 1.17 | 0.73 | 1.62 | 1.02E-03 | 8.96E-03 |
| *ADCY2* | 108 | 1.55 | 0.97 | 2.14 | 1.05E-03 | 9.04E-03 |
| *LYZL4* | 131375 | 1.04 | 0.65 | 1.43 | 1.06E-03 | 9.10E-03 |
| *LHX6* | 26468 | 1.79 | 1.11 | 2.48 | 1.07E-03 | 9.11E-03 |
| *ZNF727* | 442319 | 1.50 | 0.92 | 2.07 | 1.11E-03 | 9.25E-03 |
| *HTR2A* | 3356 | 1.38 | 0.84 | 1.91 | 1.18E-03 | 9.59E-03 |
| *PKD2L1* | 9033 | 1.49 | 0.91 | 2.07 | 1.19E-03 | 9.65E-03 |
| *DLX1* | 1745 | 1.57 | 0.96 | 2.19 | 1.25E-03 | 9.90E-03 |
| *C13orf36* | 400120 | 1.38 | 0.83 | 1.93 | 1.29E-03 | 1.01E-02 |
| *EMX2* | 2018 | 1.16 | 0.70 | 1.62 | 1.31E-03 | 1.01E-02 |
| *CXCL14* | 9547 | 1.15 | 0.69 | 1.61 | 1.35E-03 | 1.03E-02 |
| *KCNF1* | 3754 | 1.48 | 0.88 | 2.07 | 1.41E-03 | 1.05E-02 |
| *RASL10A* | 10633 | 1.12 | 0.67 | 1.57 | 1.43E-03 | 1.06E-02 |
| *LY86-AS1* | 285780 | 1.93 | 1.15 | 2.72 | 1.44E-03 | 1.06E-02 |
| *NGEF* | 25791 | 1.42 | 0.84 | 2.00 | 1.52E-03 | 1.10E-02 |
| *TM7SF4* | 81501 | 1.04 | 0.61 | 1.46 | 1.53E-03 | 1.10E-02 |
| *PPEF1* | 5475 | 1.13 | 0.67 | 1.60 | 1.55E-03 | 1.11E-02 |
| *TRIM54* | 57159 | 1.71 | 0.99 | 2.43 | 1.68E-03 | 1.16E-02 |
| *HS3ST2* | 9956 | 1.09 | 0.63 | 1.55 | 1.72E-03 | 1.18E-02 |
| *SATB2* | 23314 | 1.23 | 0.71 | 1.75 | 1.79E-03 | 1.21E-02 |
| *NECAB2* | 54550 | 1.18 | 0.68 | 1.68 | 1.80E-03 | 1.21E-02 |
| *SLC26A4* | 5172 | 1.34 | 0.76 | 1.92 | 1.91E-03 | 1.25E-02 |
| *PCSK1* | 5122 | 1.15 | 0.65 | 1.65 | 1.92E-03 | 1.25E-02 |
| *RPRML* | 388394 | 1.19 | 0.66 | 1.71 | 2.09E-03 | 1.31E-02 |
| *AC109486.1* | 25859 | 1.03 | 0.57 | 1.50 | 2.28E-03 | 1.37E-02 |
| *KCNS1* | 3787 | 1.54 | 0.84 | 2.24 | 2.35E-03 | 1.40E-02 |
| *KIAA0748* | 9840 | 1.17 | 0.63 | 1.70 | 2.47E-03 | 1.44E-02 |
| *AC002563.2* | 387890 | 1.28 | 0.69 | 1.87 | 2.55E-03 | 1.46E-02 |
| *MYB* | 4602 | 1.03 | 0.55 | 1.50 | 2.59E-03 | 1.47E-02 |
| *FREM3* | 166752 | 1.05 | 0.56 | 1.53 | 2.63E-03 | 1.49E-02 |
| *LOC646627* | 646627 | 1.73 | 0.93 | 2.53 | 2.64E-03 | 1.49E-02 |
| *AC079341.1* | 100192379 | 1.33 | 0.71 | 1.95 | 2.70E-03 | 1.51E-02 |
| *CRH* | 1392 | 1.19 | 0.63 | 1.75 | 2.78E-03 | 1.54E-02 |
| *THEMIS* | 387357 | 1.57 | 0.80 | 2.34 | 3.37E-03 | 1.73E-02 |
| *NEUROD2* | 4761 | 1.06 | 0.52 | 1.60 | 3.96E-03 | 1.91E-02 |
| *KCNJ4* | 3761 | 1.50 | 0.73 | 2.27 | 4.04E-03 | 1.93E-02 |
| *ZBBX* | 79740 | 1.06 | 0.52 | 1.60 | 4.07E-03 | 1.94E-02 |
| *GALNTL5* | 168391 | 1.40 | 0.68 | 2.13 | 4.23E-03 | 1.98E-02 |
| *ANXA8* | 653145 | 1.22 | 0.58 | 1.86 | 4.45E-03 | 2.04E-02 |
| *C6orf105* | 84830 | 1.08 | 0.51 | 1.65 | 4.65E-03 | 2.10E-02 |
| *RXFP1* | 59350 | 1.09 | 0.48 | 1.70 | 6.04E-03 | 2.48E-02 |
| *GAST* | 2520 | 1.25 | 0.53 | 1.97 | 6.46E-03 | 2.59E-02 |
| *GLP2R* | 9340 | 1.04 | 0.43 | 1.66 | 7.26E-03 | 2.79E-02 |
| *FAP* | 2191 | 1.02 | 0.38 | 1.67 | 9.73E-03 | 3.37E-02 |

Supplementary Table 5 Differentially upregulated genes in the anterior cingulate network (D).

| **Gene** | **Entrez ID** | **Fold-change** | **Lower95** | **Upper95** | ***P*-value** | **BH** |
| --- | --- | --- | --- | --- | --- | --- |
| *ADRA1B* | 147 | 1.24 | 1.04 | 1.43 | 1.69E-05 | 6.39E-03 |
| *MPPED1* | 758 | 1.81 | 1.40 | 2.22 | 9.61E-05 | 6.39E-03 |
| *CCKBR* | 887 | 1.37 | 1.09 | 1.66 | 6.08E-05 | 6.39E-03 |
| *CDH13* | 1012 | 1.42 | 1.15 | 1.70 | 4.54E-05 | 6.39E-03 |
| *CENPF* | 1063 | 1.01 | 0.82 | 1.20 | 3.89E-05 | 6.39E-03 |
| *COL5A2* | 1290 | 1.54 | 1.22 | 1.86 | 6.45E-05 | 6.39E-03 |
| *CRYM* | 1428 | 1.66 | 1.39 | 1.92 | 1.72E-05 | 6.39E-03 |
| *FHL2* | 2274 | 1.56 | 1.23 | 1.90 | 7.07E-05 | 6.39E-03 |
| *HTR1A* | 3350 | 1.06 | 0.82 | 1.31 | 9.69E-05 | 6.39E-03 |
| *KCNC2* | 3747 | 1.64 | 1.29 | 1.99 | 7.42E-05 | 6.39E-03 |
| *MATK* | 4145 | 1.07 | 0.83 | 1.31 | 8.78E-05 | 6.39E-03 |
| *MEF2C* | 4208 | 1.75 | 1.33 | 2.16 | 1.16E-04 | 6.39E-03 |
| *NEUROD2* | 4761 | 1.63 | 1.37 | 1.88 | 1.43E-05 | 6.39E-03 |
| *NPPA* | 4878 | 2.09 | 1.68 | 2.50 | 4.84E-05 | 6.39E-03 |
| *NPTX1* | 4884 | 1.17 | 0.98 | 1.36 | 1.89E-05 | 6.39E-03 |
| *NPTX2* | 4885 | 1.28 | 1.02 | 1.54 | 5.42E-05 | 6.39E-03 |
| *SERPINF1* | 5176 | 1.50 | 1.23 | 1.77 | 3.04E-05 | 6.39E-03 |
| *VIT* | 5212 | 1.03 | 0.83 | 1.23 | 4.45E-05 | 6.39E-03 |
| *PPEF1* | 5475 | 1.79 | 1.40 | 2.18 | 7.95E-05 | 6.39E-03 |
| *KLK7* | 5650 | 1.39 | 1.16 | 1.62 | 1.99E-05 | 6.39E-03 |
| *RASGRF2* | 5924 | 1.00 | 0.77 | 1.23 | 9.96E-05 | 6.39E-03 |
| *RS1* | 6247 | 1.44 | 1.16 | 1.72 | 4.17E-05 | 6.39E-03 |
| *SLN* | 6588 | 2.32 | 1.82 | 2.82 | 7.47E-05 | 6.39E-03 |
| *NR2E1* | 7101 | 1.18 | 0.90 | 1.46 | 1.14E-04 | 6.39E-03 |
| *SLC30A3* | 7781 | 2.56 | 2.10 | 3.02 | 3.03E-05 | 6.39E-03 |
| *RASAL1* | 8437 | 1.11 | 0.87 | 1.35 | 7.39E-05 | 6.39E-03 |
| *DOC2A* | 8448 | 1.16 | 0.97 | 1.36 | 1.95E-05 | 6.39E-03 |
| *HSPB3* | 8988 | 2.91 | 2.32 | 3.51 | 5.64E-05 | 6.39E-03 |
| *DLGAP2* | 9228 | 1.35 | 1.05 | 1.65 | 8.55E-05 | 6.39E-03 |
| *GLP2R* | 9340 | 1.44 | 1.11 | 1.77 | 9.47E-05 | 6.39E-03 |
| *CABP1* | 9478 | 1.69 | 1.42 | 1.97 | 1.86E-05 | 6.39E-03 |
| *MAFB* | 9935 | 1.14 | 0.97 | 1.30 | 1.13E-05 | 6.39E-03 |
| *HS3ST2* | 9956 | 2.13 | 1.68 | 2.58 | 6.32E-05 | 6.39E-03 |
| *SATB2* | 23314 | 2.09 | 1.60 | 2.58 | 1.14E-04 | 6.39E-03 |
| *LRRC8B* | 23507 | 1.03 | 0.83 | 1.24 | 5.30E-05 | 6.39E-03 |
| *SOSTDC1* | 25928 | 1.14 | 0.90 | 1.38 | 6.81E-05 | 6.39E-03 |
| *ATRNL1* | 26033 | 1.12 | 0.91 | 1.33 | 3.58E-05 | 6.39E-03 |
| *TIAM2* | 26230 | 1.17 | 0.95 | 1.39 | 3.49E-05 | 6.39E-03 |
| *AK5* | 26289 | 1.09 | 0.83 | 1.35 | 1.15E-04 | 6.39E-03 |
| *KCNV1* | 27012 | 2.25 | 1.78 | 2.73 | 6.64E-05 | 6.39E-03 |
| *RND1* | 27289 | 1.06 | 0.83 | 1.28 | 6.50E-05 | 6.39E-03 |
| *ASB2* | 51676 | 1.98 | 1.55 | 2.41 | 7.66E-05 | 6.39E-03 |
| *BCL11A* | 53335 | 1.03 | 0.79 | 1.26 | 9.34E-05 | 6.39E-03 |
| *SHC3* | 53358 | 1.13 | 0.88 | 1.38 | 8.10E-05 | 6.39E-03 |
| *FEZF2* | 55079 | 2.10 | 1.61 | 2.59 | 1.06E-04 | 6.39E-03 |
| *SLC17A7* | 57030 | 1.88 | 1.58 | 2.17 | 1.53E-05 | 6.39E-03 |
| *CAMK1G* | 57172 | 1.03 | 0.80 | 1.27 | 1.00E-04 | 6.39E-03 |
| *DPP10* | 57628 | 1.08 | 0.88 | 1.28 | 3.36E-05 | 6.39E-03 |
| *FAM5B* | 57795 | 1.12 | 0.86 | 1.38 | 1.04E-04 | 6.39E-03 |
| *RXFP1* | 59350 | 2.09 | 1.60 | 2.57 | 1.08E-04 | 6.39E-03 |
| *C6orf105* | 84830 | 2.04 | 1.59 | 2.50 | 8.43E-05 | 6.39E-03 |
| *DUSP27* | 92235 | 1.53 | 1.19 | 1.87 | 8.06E-05 | 6.39E-03 |
| *SH2D1B* | 117157 | 1.40 | 1.10 | 1.70 | 7.03E-05 | 6.39E-03 |
| *MUCL1* | 118430 | 1.24 | 1.00 | 1.49 | 4.81E-05 | 6.39E-03 |
| *RHEBL1* | 121268 | 1.29 | 1.02 | 1.57 | 6.58E-05 | 6.39E-03 |
| *C13orf16* | 121793 | 1.55 | 1.25 | 1.85 | 4.51E-05 | 6.39E-03 |
| *LYZL4* | 131375 | 1.29 | 0.99 | 1.60 | 1.08E-04 | 6.39E-03 |
| *TMEM155* | 132332 | 3.53 | 2.85 | 4.21 | 4.14E-05 | 6.39E-03 |
| *RTN4RL1* | 146760 | 1.26 | 1.02 | 1.49 | 3.69E-05 | 6.39E-03 |
| *WDR16* | 146845 | 1.12 | 0.86 | 1.38 | 9.87E-05 | 6.39E-03 |
| *ARX* | 170302 | 1.09 | 0.91 | 1.27 | 2.21E-05 | 6.39E-03 |
| *ANKRD24* | 170961 | 1.01 | 0.79 | 1.23 | 8.32E-05 | 6.39E-03 |
| *DHRS7C* | 201140 | 1.27 | 1.02 | 1.52 | 4.53E-05 | 6.39E-03 |
| *LY86-AS1* | 285780 | 2.80 | 2.30 | 3.30 | 2.98E-05 | 6.39E-03 |
| *FAM19A2* | 338811 | 1.35 | 1.05 | 1.65 | 8.78E-05 | 6.39E-03 |
| *LOC339524* | 339524 | 1.40 | 1.15 | 1.64 | 2.69E-05 | 6.39E-03 |
| *C2orf55* | 343990 | 1.52 | 1.16 | 1.87 | 1.09E-04 | 6.39E-03 |
| *LCE3C* | 353144 | 1.01 | 0.80 | 1.22 | 6.51E-05 | 6.39E-03 |
| *THEMIS* | 387357 | 2.83 | 2.17 | 3.50 | 1.10E-04 | 6.39E-03 |
| *AC002563.2* | 387890 | 2.28 | 1.95 | 2.62 | 1.17E-05 | 6.39E-03 |
| *C6orf126* | 389383 | 1.37 | 1.09 | 1.66 | 6.05E-05 | 6.39E-03 |
| *C1QL1* | 389941 | 1.83 | 1.44 | 2.23 | 7.28E-05 | 6.39E-03 |
| *KRT16P2* | 400578 | 1.48 | 1.21 | 1.74 | 2.93E-05 | 6.39E-03 |
| *AC112641.2* | 401097 | 1.59 | 1.27 | 1.91 | 5.43E-05 | 6.39E-03 |
| *ZNF727* | 442319 | 1.80 | 1.43 | 2.18 | 6.06E-05 | 6.39E-03 |
| *LOC643750* | 643750 | 1.01 | 0.85 | 1.18 | 1.91E-05 | 6.39E-03 |
| *KCNS1* | 3787 | 2.59 | 1.97 | 3.22 | 1.22E-04 | 6.44E-03 |
| *EPHX4* | 253152 | 1.02 | 0.78 | 1.27 | 1.22E-04 | 6.44E-03 |
| *LOC100287347* | 100287347 | 1.15 | 0.88 | 1.43 | 1.21E-04 | 6.44E-03 |
| *CCK* | 885 | 3.00 | 2.27 | 3.73 | 1.29E-04 | 6.46E-03 |
| *LHX2* | 9355 | 2.00 | 1.52 | 2.48 | 1.26E-04 | 6.46E-03 |
| *GDA* | 9615 | 2.83 | 2.15 | 3.52 | 1.28E-04 | 6.46E-03 |
| *AC078937.4* | 286002 | 2.28 | 1.73 | 2.83 | 1.26E-04 | 6.46E-03 |
| *CASQ1* | 844 | 1.53 | 1.16 | 1.90 | 1.30E-04 | 6.46E-03 |
| *KIAA0748* | 9840 | 2.44 | 1.85 | 3.03 | 1.30E-04 | 6.46E-03 |
| *TNNT2* | 7139 | 2.35 | 1.78 | 2.92 | 1.32E-04 | 6.47E-03 |
| *CREG2* | 200407 | 2.07 | 1.57 | 2.58 | 1.36E-04 | 6.54E-03 |
| *DACH2* | 117154 | 1.03 | 0.77 | 1.28 | 1.41E-04 | 6.55E-03 |
| *SLC26A4* | 5172 | 1.83 | 1.38 | 2.29 | 1.42E-04 | 6.56E-03 |
| *CDH9* | 1007 | 1.47 | 1.11 | 1.84 | 1.45E-04 | 6.64E-03 |
| *CYP26A1* | 1592 | 1.69 | 1.27 | 2.11 | 1.50E-04 | 6.64E-03 |
| *RASL11B* | 65997 | 1.20 | 0.90 | 1.50 | 1.50E-04 | 6.64E-03 |
| *SGK493* | 91461 | 1.31 | 0.98 | 1.63 | 1.52E-04 | 6.69E-03 |
| *NPFFR2* | 10886 | 1.24 | 0.93 | 1.55 | 1.53E-04 | 6.70E-03 |
| *ADCY2* | 108 | 1.91 | 1.42 | 2.39 | 1.61E-04 | 6.76E-03 |
| *EPHB6* | 2051 | 1.45 | 1.09 | 1.82 | 1.57E-04 | 6.76E-03 |
| *EXTL1* | 2134 | 1.15 | 0.86 | 1.44 | 1.64E-04 | 6.76E-03 |
| *LMO4* | 8543 | 1.37 | 1.02 | 1.72 | 1.62E-04 | 6.76E-03 |
| *KCNH3* | 23416 | 1.10 | 0.82 | 1.38 | 1.59E-04 | 6.76E-03 |
| *OVOL2* | 58495 | 1.70 | 1.27 | 2.13 | 1.58E-04 | 6.76E-03 |
| *NEUROD6* | 63974 | 1.92 | 1.43 | 2.40 | 1.58E-04 | 6.76E-03 |
| *OSBPL3* | 26031 | 1.22 | 0.91 | 1.54 | 1.66E-04 | 6.80E-03 |
| *NNMT* | 4837 | 1.23 | 0.91 | 1.54 | 1.67E-04 | 6.81E-03 |
| *RGS4* | 5999 | 1.92 | 1.43 | 2.42 | 1.71E-04 | 6.83E-03 |
| *AC010087.3* | 129293 | 1.40 | 1.04 | 1.76 | 1.76E-04 | 6.89E-03 |
| *RSPO2* | 340419 | 1.73 | 1.28 | 2.18 | 1.80E-04 | 6.89E-03 |
| *VIP* | 7432 | 1.85 | 1.36 | 2.33 | 1.83E-04 | 6.89E-03 |
| *VSNL1* | 7447 | 1.01 | 0.74 | 1.27 | 1.84E-04 | 6.89E-03 |
| *NMU* | 10874 | 1.47 | 1.09 | 1.85 | 1.85E-04 | 6.89E-03 |
| *AC079341.1* | 100192379 | 2.18 | 1.61 | 2.74 | 1.83E-04 | 6.89E-03 |
| *OLFM1* | 10439 | 1.04 | 0.76 | 1.31 | 1.89E-04 | 6.90E-03 |
| *ST6GALNAC5* | 81849 | 1.21 | 0.89 | 1.53 | 1.89E-04 | 6.90E-03 |
| *CRYBB1* | 1414 | 1.09 | 0.80 | 1.37 | 1.91E-04 | 6.90E-03 |
| *C6orf142* | 90523 | 1.83 | 1.34 | 2.31 | 1.94E-04 | 6.90E-03 |
| *KCNS2* | 3788 | 1.29 | 0.95 | 1.63 | 1.94E-04 | 6.91E-03 |
| *RBP4* | 5950 | 1.50 | 1.10 | 1.90 | 1.96E-04 | 6.91E-03 |
| *MCHR2* | 84539 | 1.71 | 1.26 | 2.16 | 1.95E-04 | 6.91E-03 |
| *MUM1L1* | 139221 | 1.53 | 1.12 | 1.94 | 2.00E-04 | 6.91E-03 |
| *GALNTL5* | 168391 | 2.08 | 1.53 | 2.64 | 2.00E-04 | 6.91E-03 |
| *DLX6-AS1* | 285987 | 1.32 | 0.97 | 1.67 | 1.98E-04 | 6.91E-03 |
| *PTGS2* | 5743 | 1.25 | 0.92 | 1.59 | 2.10E-04 | 6.93E-03 |
| *LMO3* | 55885 | 1.17 | 0.86 | 1.49 | 2.10E-04 | 6.93E-03 |
| *FAM162B* | 221303 | 1.10 | 0.80 | 1.39 | 2.07E-04 | 6.93E-03 |
| *C2orf80* | 389073 | 1.10 | 0.81 | 1.40 | 2.09E-04 | 6.93E-03 |
| *SOHLH1* | 402381 | 1.53 | 1.12 | 1.94 | 2.06E-04 | 6.93E-03 |
| *ENC1* | 8507 | 1.83 | 1.34 | 2.33 | 2.21E-04 | 7.11E-03 |
| *MAPK13* | 5603 | 1.00 | 0.73 | 1.28 | 2.22E-04 | 7.11E-03 |
| *C17orf96* | 100170841 | 1.55 | 1.13 | 1.97 | 2.22E-04 | 7.11E-03 |
| *AQP9* | 366 | 1.47 | 1.07 | 1.87 | 2.24E-04 | 7.13E-03 |
| *CHN1* | 1123 | 1.13 | 0.82 | 1.43 | 2.31E-04 | 7.13E-03 |
| *FOXG1B* | 2290 | 2.77 | 2.01 | 3.53 | 2.29E-04 | 7.13E-03 |
| *CACNG3* | 10368 | 1.55 | 1.13 | 1.97 | 2.25E-04 | 7.13E-03 |
| *POU6F2* | 11281 | 1.09 | 0.79 | 1.39 | 2.28E-04 | 7.13E-03 |
| *ABCC12* | 94160 | 1.77 | 1.28 | 2.25 | 2.30E-04 | 7.13E-03 |
| *PKD2L1* | 9033 | 2.11 | 1.53 | 2.69 | 2.35E-04 | 7.19E-03 |
| *GABRA5* | 2558 | 1.41 | 1.02 | 1.79 | 2.36E-04 | 7.19E-03 |
| *MYO5B* | 4645 | 1.10 | 0.79 | 1.40 | 2.40E-04 | 7.21E-03 |
| *MCHR1* | 2847 | 1.29 | 0.93 | 1.65 | 2.44E-04 | 7.22E-03 |
| *AC109486.1* | 25859 | 2.01 | 1.45 | 2.57 | 2.47E-04 | 7.26E-03 |
| *GPR26* | 2849 | 1.82 | 1.31 | 2.33 | 2.50E-04 | 7.27E-03 |
| *LDB2* | 9079 | 1.43 | 1.03 | 1.83 | 2.51E-04 | 7.27E-03 |
| *OR14I1* | 401994 | 2.08 | 1.50 | 2.66 | 2.57E-04 | 7.31E-03 |
| *HTR2A* | 3356 | 1.94 | 1.39 | 2.48 | 2.62E-04 | 7.35E-03 |
| *MOXD1* | 26002 | 1.57 | 1.12 | 2.01 | 2.68E-04 | 7.39E-03 |
| *BAIAP3* | 8938 | 1.20 | 0.86 | 1.54 | 2.72E-04 | 7.42E-03 |
| *NRGN* | 4900 | 2.61 | 1.87 | 3.36 | 2.76E-04 | 7.44E-03 |
| *RPRM* | 56475 | 1.29 | 0.92 | 1.66 | 2.77E-04 | 7.44E-03 |
| *STX1A* | 6804 | 1.42 | 1.01 | 1.82 | 2.82E-04 | 7.46E-03 |
| *EGR3* | 1960 | 1.69 | 1.20 | 2.17 | 2.88E-04 | 7.52E-03 |
| *DLX1* | 1745 | 1.91 | 1.36 | 2.46 | 2.91E-04 | 7.54E-03 |
| *TMEM132D* | 121256 | 1.26 | 0.90 | 1.62 | 2.94E-04 | 7.55E-03 |
| *PCDH8* | 5100 | 1.30 | 0.93 | 1.68 | 2.99E-04 | 7.58E-03 |
| *TBR1* | 10716 | 1.59 | 1.13 | 2.05 | 3.07E-04 | 7.62E-03 |
| *NUDT4P1* | 11163 | 1.19 | 0.85 | 1.54 | 3.09E-04 | 7.63E-03 |
| *KIAA1239* | 57495 | 1.43 | 1.02 | 1.85 | 3.14E-04 | 7.65E-03 |
| *NEK2* | 4751 | 1.23 | 0.87 | 1.59 | 3.29E-04 | 7.74E-03 |
| *KCNT2* | 343450 | 1.02 | 0.72 | 1.33 | 3.40E-04 | 7.85E-03 |
| *CRHBP* | 1393 | 1.26 | 0.88 | 1.63 | 3.43E-04 | 7.87E-03 |
| *ITPKA* | 3706 | 1.48 | 1.04 | 1.92 | 3.51E-04 | 7.93E-03 |
| *DNAJC5G* | 285126 | 1.02 | 0.72 | 1.33 | 3.53E-04 | 7.95E-03 |
| *WIF1* | 11197 | 1.03 | 0.72 | 1.34 | 3.55E-04 | 7.95E-03 |
| *KCNJ4* | 3761 | 1.73 | 1.21 | 2.25 | 3.61E-04 | 7.95E-03 |
| *AC005551.1* | 284422 | 1.10 | 0.77 | 1.43 | 3.66E-04 | 7.96E-03 |
| *HRH1* | 3269 | 1.56 | 1.09 | 2.04 | 3.69E-04 | 7.99E-03 |
| *PDE2A* | 5138 | 1.30 | 0.90 | 1.69 | 3.76E-04 | 8.08E-03 |
| *GUCA1B* | 2979 | 1.15 | 0.80 | 1.50 | 3.81E-04 | 8.15E-03 |
| *STYK1* | 55359 | 1.19 | 0.83 | 1.55 | 3.82E-04 | 8.15E-03 |
| *CDKL1* | 8814 | 1.00 | 0.69 | 1.31 | 4.15E-04 | 8.35E-03 |
| *KMO* | 8564 | 1.26 | 0.87 | 1.65 | 4.22E-04 | 8.41E-03 |
| *RASL10A* | 10633 | 1.30 | 0.90 | 1.71 | 4.31E-04 | 8.45E-03 |
| *CAMK2A* | 815 | 1.61 | 1.10 | 2.12 | 4.72E-04 | 8.72E-03 |
| *DDN* | 23109 | 1.21 | 0.82 | 1.60 | 4.84E-04 | 8.79E-03 |
| *ERICH1-AS1* | 619343 | 1.06 | 0.72 | 1.39 | 4.87E-04 | 8.83E-03 |
| *GYPE* | 2996 | 1.24 | 0.84 | 1.64 | 4.91E-04 | 8.87E-03 |
| *LOC100290023* | 100290023 | 1.45 | 0.98 | 1.91 | 4.92E-04 | 8.87E-03 |
| *MEPE* | 56955 | 1.07 | 0.73 | 1.42 | 5.04E-04 | 8.96E-03 |
| *SLCO2A1* | 6578 | 1.11 | 0.75 | 1.47 | 5.12E-04 | 8.99E-03 |
| *KCNQ5* | 56479 | 1.03 | 0.69 | 1.36 | 5.16E-04 | 8.99E-03 |
| *AC116165.2* | 283767 | 1.24 | 0.84 | 1.65 | 5.15E-04 | 8.99E-03 |
| *SCARA5* | 286133 | 1.34 | 0.91 | 1.78 | 5.16E-04 | 8.99E-03 |
| *FAM148C* | 126567 | 1.22 | 0.82 | 1.61 | 5.17E-04 | 8.99E-03 |
| *SLIT1* | 6585 | 1.28 | 0.86 | 1.70 | 5.33E-04 | 9.06E-03 |
| *MEIS3P2* | 56917 | 1.23 | 0.83 | 1.63 | 5.41E-04 | 9.08E-03 |
| *LRRTM4* | 80059 | 1.14 | 0.77 | 1.52 | 5.46E-04 | 9.12E-03 |
| *LY6H* | 4062 | 1.10 | 0.74 | 1.47 | 5.50E-04 | 9.14E-03 |
| *CHRM3* | 1131 | 1.57 | 1.05 | 2.09 | 5.54E-04 | 9.19E-03 |
| *TOX* | 9760 | 1.03 | 0.69 | 1.37 | 5.60E-04 | 9.25E-03 |
| *CRLF1* | 9244 | 1.16 | 0.77 | 1.54 | 5.65E-04 | 9.27E-03 |
| *FMN1* | 342184 | 1.15 | 0.77 | 1.53 | 5.76E-04 | 9.32E-03 |
| *NETO1* | 81832 | 1.33 | 0.89 | 1.78 | 5.82E-04 | 9.36E-03 |
| *ZBBX* | 79740 | 1.40 | 0.93 | 1.87 | 6.00E-04 | 9.52E-03 |
| *FREM3* | 166752 | 2.22 | 1.48 | 2.96 | 6.03E-04 | 9.53E-03 |
| *B3GALT2* | 8707 | 1.01 | 0.67 | 1.35 | 6.07E-04 | 9.54E-03 |
| *ANKRD56* | 345079 | 1.77 | 1.17 | 2.36 | 6.09E-04 | 9.57E-03 |
| *KHDRBS2* | 202559 | 1.37 | 0.91 | 1.83 | 6.14E-04 | 9.60E-03 |
| *C13orf39* | 196541 | 1.05 | 0.70 | 1.40 | 6.16E-04 | 9.61E-03 |
| *TYRP1* | 7306 | 1.43 | 0.94 | 1.92 | 6.39E-04 | 9.82E-03 |
| *TRIM54* | 57159 | 2.21 | 1.46 | 2.96 | 6.48E-04 | 9.89E-03 |
| *HGF* | 3082 | 1.29 | 0.85 | 1.73 | 6.77E-04 | 9.93E-03 |
| *NPY* | 4852 | 1.86 | 1.22 | 2.50 | 6.59E-04 | 9.93E-03 |
| *ICAM5* | 7087 | 1.52 | 0.99 | 2.04 | 6.76E-04 | 9.93E-03 |
| *NTN4* | 59277 | 1.05 | 0.69 | 1.42 | 6.64E-04 | 9.93E-03 |
| *CRH* | 1392 | 1.50 | 0.98 | 2.01 | 6.80E-04 | 9.94E-03 |
| *BAIAP2L2* | 80115 | 1.12 | 0.74 | 1.51 | 6.80E-04 | 9.94E-03 |
| *FAP* | 2191 | 1.16 | 0.76 | 1.57 | 6.93E-04 | 9.99E-03 |
| *C14orf23* | 387978 | 1.18 | 0.77 | 1.59 | 6.93E-04 | 9.99E-03 |
| *KCNF1* | 3754 | 1.54 | 1.00 | 2.07 | 7.06E-04 | 1.01E-02 |
| *COL24A1* | 255631 | 1.15 | 0.75 | 1.55 | 7.14E-04 | 1.01E-02 |
| *TAC3* | 6866 | 1.83 | 1.19 | 2.47 | 7.26E-04 | 1.02E-02 |
| *ANO3* | 63982 | 1.23 | 0.80 | 1.66 | 7.38E-04 | 1.02E-02 |
| *THEM5* | 284486 | 1.05 | 0.67 | 1.42 | 7.80E-04 | 1.04E-02 |
| *PDZRN3* | 23024 | 1.19 | 0.77 | 1.62 | 7.84E-04 | 1.05E-02 |
| *CXCL14* | 9547 | 1.08 | 0.70 | 1.47 | 7.95E-04 | 1.05E-02 |
| *NGEF* | 25791 | 1.66 | 1.06 | 2.25 | 8.13E-04 | 1.06E-02 |
| *DLX2* | 1746 | 1.21 | 0.78 | 1.64 | 8.16E-04 | 1.06E-02 |
| *SLC22A9* | 114571 | 1.27 | 0.81 | 1.73 | 8.30E-04 | 1.06E-02 |
| *TC2N* | 123036 | 1.11 | 0.71 | 1.51 | 8.22E-04 | 1.06E-02 |
| *PCSK1* | 5122 | 1.57 | 1.00 | 2.13 | 8.42E-04 | 1.06E-02 |
| *C1orf115* | 79762 | 1.27 | 0.81 | 1.73 | 8.39E-04 | 1.06E-02 |
| *ZBTB16* | 7704 | 1.01 | 0.65 | 1.38 | 8.58E-04 | 1.07E-02 |
| *OPN3* | 23596 | 1.25 | 0.80 | 1.71 | 8.61E-04 | 1.07E-02 |
| *FAM19A1* | 407738 | 1.25 | 0.80 | 1.71 | 8.65E-04 | 1.07E-02 |
| *KIF17* | 57576 | 1.16 | 0.74 | 1.58 | 8.87E-04 | 1.09E-02 |
| *LOC646627* | 646627 | 2.04 | 1.29 | 2.78 | 9.04E-04 | 1.10E-02 |
| *STEAP1* | 26872 | 1.11 | 0.71 | 1.52 | 9.08E-04 | 1.10E-02 |
| *ANXA8* | 653145 | 1.71 | 1.08 | 2.34 | 9.18E-04 | 1.10E-02 |
| *GAST* | 2520 | 1.22 | 0.77 | 1.67 | 9.42E-04 | 1.11E-02 |
| *SSTR1* | 6751 | 1.04 | 0.66 | 1.42 | 9.48E-04 | 1.12E-02 |
| *LHX6* | 26468 | 1.80 | 1.13 | 2.47 | 9.74E-04 | 1.13E-02 |
| *PRSS16* | 10279 | 1.13 | 0.71 | 1.55 | 9.89E-04 | 1.13E-02 |
| *RPRML* | 388394 | 1.52 | 0.95 | 2.09 | 9.87E-04 | 1.13E-02 |
| *PHLDA2* | 7262 | 1.02 | 0.64 | 1.41 | 1.02E-03 | 1.15E-02 |
| *C13orf36* | 400120 | 1.55 | 0.96 | 2.13 | 1.05E-03 | 1.17E-02 |
| *AC017096.1* | 150538 | 1.50 | 0.93 | 2.07 | 1.07E-03 | 1.18E-02 |
| *ANKRD62* | 342850 | 1.02 | 0.63 | 1.42 | 1.09E-03 | 1.20E-02 |
| *CORT* | 1325 | 1.10 | 0.68 | 1.51 | 1.10E-03 | 1.20E-02 |
| *CIDEA* | 1149 | 1.13 | 0.69 | 1.57 | 1.17E-03 | 1.23E-02 |
| *RTP1* | 132112 | 1.81 | 1.11 | 2.52 | 1.18E-03 | 1.23E-02 |
| *FAM81A* | 145773 | 1.34 | 0.82 | 1.86 | 1.20E-03 | 1.23E-02 |
| *NOS2* | 4843 | 1.01 | 0.62 | 1.41 | 1.22E-03 | 1.25E-02 |
| *EMX2* | 2018 | 1.07 | 0.65 | 1.49 | 1.25E-03 | 1.26E-02 |
| *ASGR2* | 433 | 1.12 | 0.68 | 1.57 | 1.31E-03 | 1.30E-02 |
| *IL12RB2* | 3595 | 1.04 | 0.62 | 1.45 | 1.33E-03 | 1.31E-02 |
| *SST* | 6750 | 1.60 | 0.97 | 2.24 | 1.33E-03 | 1.31E-02 |
| *FRMPD2* | 143162 | 1.60 | 0.96 | 2.24 | 1.34E-03 | 1.31E-02 |
| *PCDH20* | 64881 | 1.06 | 0.64 | 1.49 | 1.34E-03 | 1.31E-02 |
| *C8orf4* | 56892 | 1.01 | 0.60 | 1.41 | 1.42E-03 | 1.35E-02 |
| *C21orf128* | 150147 | 1.46 | 0.87 | 2.06 | 1.41E-03 | 1.35E-02 |
| *CAMKV* | 79012 | 1.18 | 0.70 | 1.65 | 1.44E-03 | 1.35E-02 |
| *GSG1* | 83445 | 1.38 | 0.81 | 1.94 | 1.51E-03 | 1.38E-02 |
| *GFRA2* | 2675 | 1.13 | 0.67 | 1.59 | 1.53E-03 | 1.39E-02 |
| *RORB* | 6096 | 1.19 | 0.70 | 1.67 | 1.53E-03 | 1.39E-02 |
| *EMX2OS* | 196047 | 1.19 | 0.70 | 1.69 | 1.53E-03 | 1.39E-02 |
| *HTR1F* | 3355 | 1.09 | 0.64 | 1.54 | 1.61E-03 | 1.42E-02 |
| *KCNH5* | 27133 | 1.34 | 0.79 | 1.90 | 1.61E-03 | 1.42E-02 |
| *CCDC3* | 83643 | 1.05 | 0.61 | 1.48 | 1.63E-03 | 1.43E-02 |
| *ZNF831* | 128611 | 1.08 | 0.62 | 1.53 | 1.76E-03 | 1.49E-02 |
| *GRASP* | 160622 | 1.09 | 0.63 | 1.56 | 1.77E-03 | 1.49E-02 |
| *ARC* | 23237 | 1.13 | 0.65 | 1.61 | 1.78E-03 | 1.50E-02 |
| *LZTS1* | 11178 | 1.08 | 0.62 | 1.54 | 1.85E-03 | 1.53E-02 |
| *AKAP5* | 9495 | 1.05 | 0.60 | 1.51 | 1.86E-03 | 1.53E-02 |
| *PAH* | 5053 | 1.19 | 0.67 | 1.71 | 1.97E-03 | 1.58E-02 |
| *GRHL2* | 79977 | 1.15 | 0.63 | 1.67 | 2.40E-03 | 1.75E-02 |
| *IGFBP2* | 3485 | 1.18 | 0.64 | 1.72 | 2.50E-03 | 1.78E-02 |
| *NCALD* | 83988 | 1.11 | 0.60 | 1.62 | 2.60E-03 | 1.81E-02 |
| *CBLN4* | 140689 | 1.03 | 0.55 | 1.51 | 2.63E-03 | 1.82E-02 |
| *FBXO40* | 51725 | 1.42 | 0.69 | 2.14 | 3.95E-03 | 2.28E-02 |
| *KCTD16* | 57528 | 1.16 | 0.56 | 1.75 | 4.10E-03 | 2.34E-02 |
| *IQGAP3* | 128239 | 1.35 | 0.64 | 2.07 | 4.51E-03 | 2.46E-02 |

Supplementary Table 6 Pathway analysis of DEGs upregulated in the posterior cingulate network (C) and anterior cingulate network (D). Many pathways are shared between both networks (red text).

| **Pathway** | **BH** | **Gene count** |
| --- | --- | --- |
| **Posterior cingulate network (C)** |  |  |
| Neuronal System | 3.19E-09 | 23 |
| Voltage gated Potassium channels | 1.44E-06 | 8 |
| Transmission across Chemical Synapses | 6.95E-06 | 15 |
| Neurotransmitter receptors and postsynaptic signal transmission | 8.32E-06 | 13 |
| GPCR ligand binding | 2.16E-05 | 18 |
| Class A/1 (Rhodopsin-like receptors) | 2.42E-05 | 15 |
| Potassium Channels | 2.65E-05 | 9 |
| Transcriptional Regulation by *MECP2* | 1.45E-03 | 6 |
| Amine ligand-binding receptors | 1.93E-03 | 5 |
| Ras activation upon Ca2+ influx through NMDA receptor | 2.03E-03 | 4 |
| Long-term potentiation | 3.12E-03 | 4 |
| CREB1 phosphorylation through NMDA receptor-mediated activation of RAS signaling | 5.96E-03 | 4 |
| Activation of NMDA receptors and postsynaptic events | 7.18E-03 | 6 |
| G alpha (q) signalling events | 7.26E-03 | 9 |
| Lysosphingolipid and LPA receptors | 7.30E-03 | 3 |
| Assembly and cell surface presentation of NMDA receptors | 1.13E-02 | 4 |
| Post NMDA receptor activation events | 2.15E-02 | 5 |
| Unblocking of NMDA receptors, glutamate binding and activation | 2.62E-02 | 3 |
| Negative regulation of NMDA receptor-mediated neuronal transmission | 2.62E-02 | 3 |
| G alpha (i) signalling events | 2.86E-02 | 11 |
| GABA receptor activation | 3.67E-02 | 4 |
| Peptide ligand-binding receptors | 3.67E-02 | 7 |
| Trafficking of AMPA receptors | 4.97E-02 | 3 |
| Glutamate binding, activation of AMPA receptors and synaptic plasticity | 4.97E-02 | 3 |
| **Anterior cingulate network (D)** |  |  |
| GPCR ligand binding | 3.34E-06 | 23 |
| Class A/1 (Rhodopsin-like receptors) | 3.34E-06 | 19 |
| Neuronal System | 6.21E-06 | 21 |
| Voltage gated Potassium channels | 6.21E-06 | 8 |
| G alpha (q) signalling events | 2.61E-04 | 13 |
| Peptide ligand-binding receptors | 2.61E-04 | 12 |
| Potassium Channels | 2.61E-04 | 9 |
| Amine ligand-binding receptors | 6.30E-04 | 6 |
| G alpha (i) signalling events | 1.73E-03 | 16 |
| Transmission across Chemical Synapses | 6.93E-03 | 12 |
| Neurotransmitter receptors and postsynaptic signal transmission | 9.83E-03 | 10 |
| Serotonin receptors | 1.35E-02 | 3 |
| Transcriptional Regulation by *MECP2* | 3.56E-02 | 5 |

Supplementary Table 7 Region-specific acronyms used as column names in heatmaps.

| **Acronym** | **Name** | **Acronym** | **Name** |
| --- | --- | --- | --- |
| AOrG | Anterior orbital gyrus | SMG-s | Supramarginal gyrus, superior bank of gyrus |
| fro | Frontal operculum | SMG-i | Supramarginal gyrus, inferior bank of gyrus |
| FP-s | Frontal pole, superior aspect | PCLp-cs | Paracentral lobule, posterior part, bank of cingulate sulcus |
| FPi | Frontal pole, inferior aspect | PoG-cs | Postcentral gyrus, bank of the central sulcus |
| GRe | Gyrus rectus | PoG-sl | Postcentral gyrus, superior lateral aspect of gyrus |
| opIFG | Inferior frontal gyrus, opercular part | PoG-il | Postcentral gyrus, inferior lateral aspect of gyrus |
| orIFG | Inferior frontal gyrus, orbital part | PoG-pcs | Postcentral gyrus, bank of the posterior central sulcus |
| trIFG | Inferior frontal gyrus, triangular part | Pcu-s | Precuneus, superior lateral bank of gyrus |
| IRoG | Inferior rostral gyrus | Pcu-i | Precuneus, inferior lateral bank of gyrus |
| LOrG | Lateral orbital gyrus | FuG-its | Fusiform gyrus, bank of the its |
| MOrG | Medial orbital gyrus | FuG-l | Fusiform gyrus, lateral bank of gyrus |
| MFG-s | Middle frontal gyrus, superior bank of gyrus | FuG-cos | Fusiform gyrus, bank of cos |
| MFG-i | Middle frontal gyrus, inferior bank of gyrus | HG | Heschl's gyrus |
| PCLa | Paracentral lobule, anterior part | ITG-its | Inferior temporal gyrus, bank of the its |
| PCLa-s | Paracentral lobule, anterior part, superior bank of gyrus | ITG-l | Inferior temporal gyrus, lateral bank of gyrus |
| PCLa-i | Paracentral lobule, anterior part, inferior bank of gyrus | ITG-mts | Inferior temporal gyrus, bank of mts |
| PaOG | Parolfactory gyri | MTG-s | Middle temporal gyrus, superior bank of gyrus |
| POrG | Posterior orbital gyrus | MTG-i | Middle temporal gyrus, inferior bank of gyrus |
| PrG-prc | Precentral gyrus, bank of the precentral sulcus | PLP | Planum polare |
| PrG-sl | Precentral gyrus, superior lateral aspect of gyrus | PLT | Planum temporale |
| PrG-il | Precentral gyrus, inferior lateral aspect of gyrus | STG-l | Superior temporal gyrus, lateral bank of gyrus |
| PrG-cs | Precentral gyrus, bank of the central sulcus | STG-i | Superior temporal gyrus, inferior bank of gyrus |
| SFG-m | Superior frontal gyrus, medial bank of gyrus | TP-s | Temporal pole, superior aspect |
| SFG-l | Superior frontal gyrus, lateral bank of gyrus | TG | Transverse gyri |
| SRoG | Superior rostral gyrus | ATZ | Amygdalohippocampal transition zone |
| LIG | Long insular gyri | BLA | Basolateral nucleus |
| SIG | Short insular gyri | BMA | Basomedial nucleus |
| CgGf-s | Cingulate gyrus, frontal part, superior bank of gyrus | CeA | Central nucleus |
| CgGf-i | Cingulate gyrus, frontal part, inferior bank of gyrus | COMA | Cortico-medial group |
| CgGp-s | Cingulate gyrus, parietal part, superior bank of gyrus | LA | Lateral nucleus |
| CgGp-i | Cingulate gyrus, parietal part, inferior bank of gyrus | DBv | Nucleus of the diagonal band, horizontal division |
| CgGr-s | Cingulate gyrus, retrosplenial part, superior bank of gyrus | OlfT | Olfactory tubercle |
| CgGr-i | Cingulate gyrus, retrosplenial part, inferior bank of gyrus | GPe | Globus pallidus, external segment |
| SCG | Subcallosal cingulate gyrus | TCd | Tail of caudate nucleus |
| DG | Dentate gyrus | Pu | Putamen |
| CA1 | CA1 field | Cl | Claustrum |
| CA2 | CA2 field | SO | Supraoptic nucleus |
| CA3 | CA3 field | Sb | Subthalamic nucleus |
| CA4 | CA4 field | LGd | Dorsal lateral geniculate nucleus |
| S | Subiculum | DTLv | Lateral group of nuclei, ventral division |
| PHG-l | Parahippocampal gyrus, lateral bank of gyrus | MG | Medial geniculate complex |
| PHG-cos | Parahippocampal gyrus, bank of the cos | DTP | Posterior group of nuclei |
| Pir | Piriform cortex | R | Reticular nucleus of thalamus |
| Cun-pest | Cuneus, peristriate | SNC | Substantia nigra, pars compacta |
| IOG-s | Inferior occipital gyrus, superior bank of gyrus | SNR | Substantia nigra, pars reticulata |
| IOG-i | Inferior occipital gyrus, inferior bank of gyrus | He-Crus I | Crus I, lateral hemisphere |
| LiG-str | Lingual gyrus, striate | PV-Crus II | Crus II, paravermis |
| OTG-s | Occipito-temporal gyrus, superior bank of gyrus | He-Crus II | Crus II, lateral hemisphere |
| OTG-i | Occipito-temporal gyrus, inferior bank of gyrus | PV-VIIB | VIIB, paravermis |
| SOG-s | Superior occipital gyrus, superior bank of gyrus | He-VIIB | VIIB, lateral hemisphere |
| SOG-i | Superior occipital gyrus, inferior bank of gyrus | He-VIIIA | VIIIA, lateral hemisphere |
| AnG-i | Angular gyrus, inferior bank of gyrus | CPLV | Choroid plexus of the lateral ventricle |

### Supplementary Figures

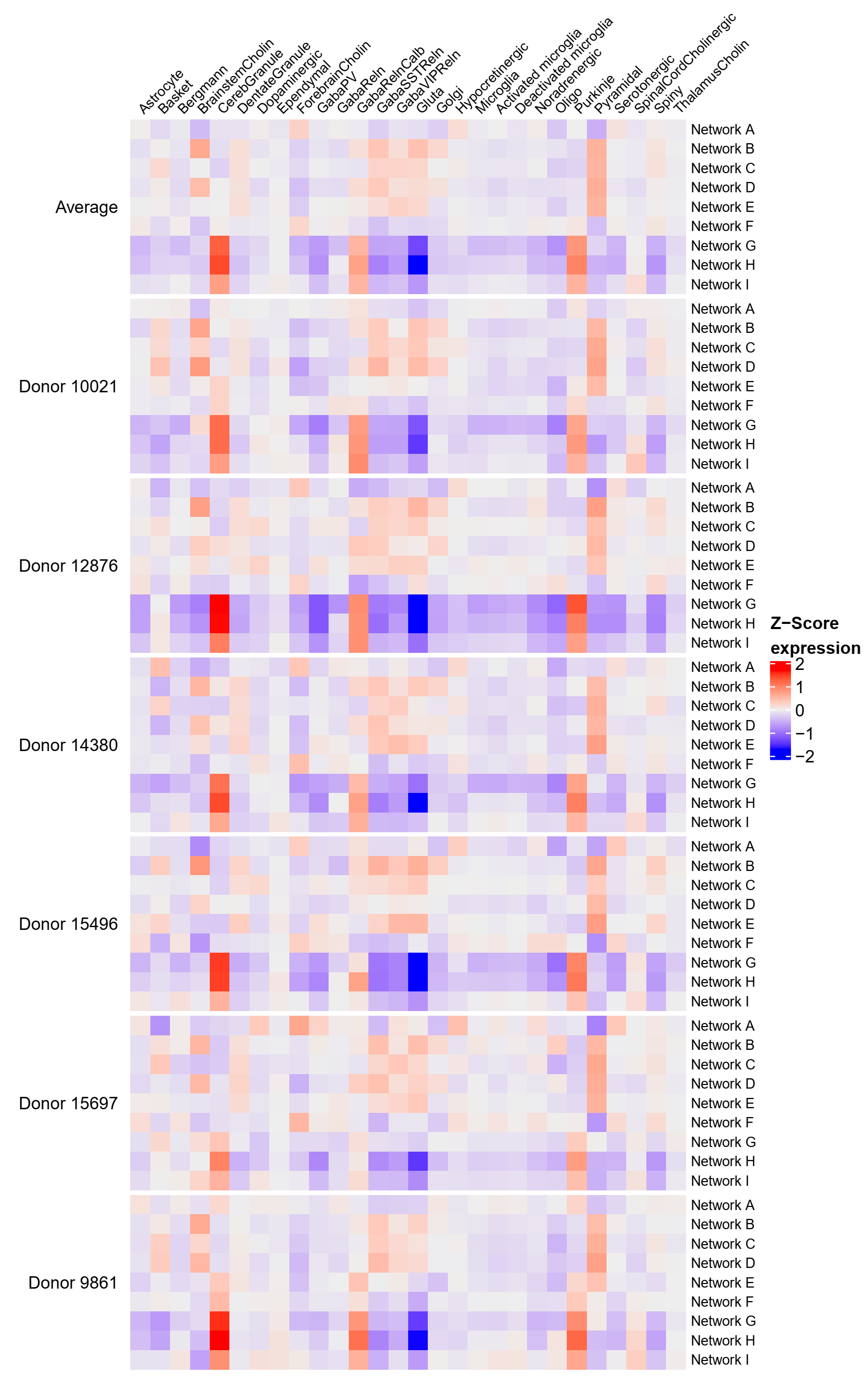

Supplementary Figure 1 Expression of cell-types in anatomical networks. Gene expression was Z-scored and averaged across cell-type specific markers and across samples within anatomical networks. First heatmap shows the element-wise average expression across the six donors in the Allen Human Brain Atlas for which these heatmaps are also shown.

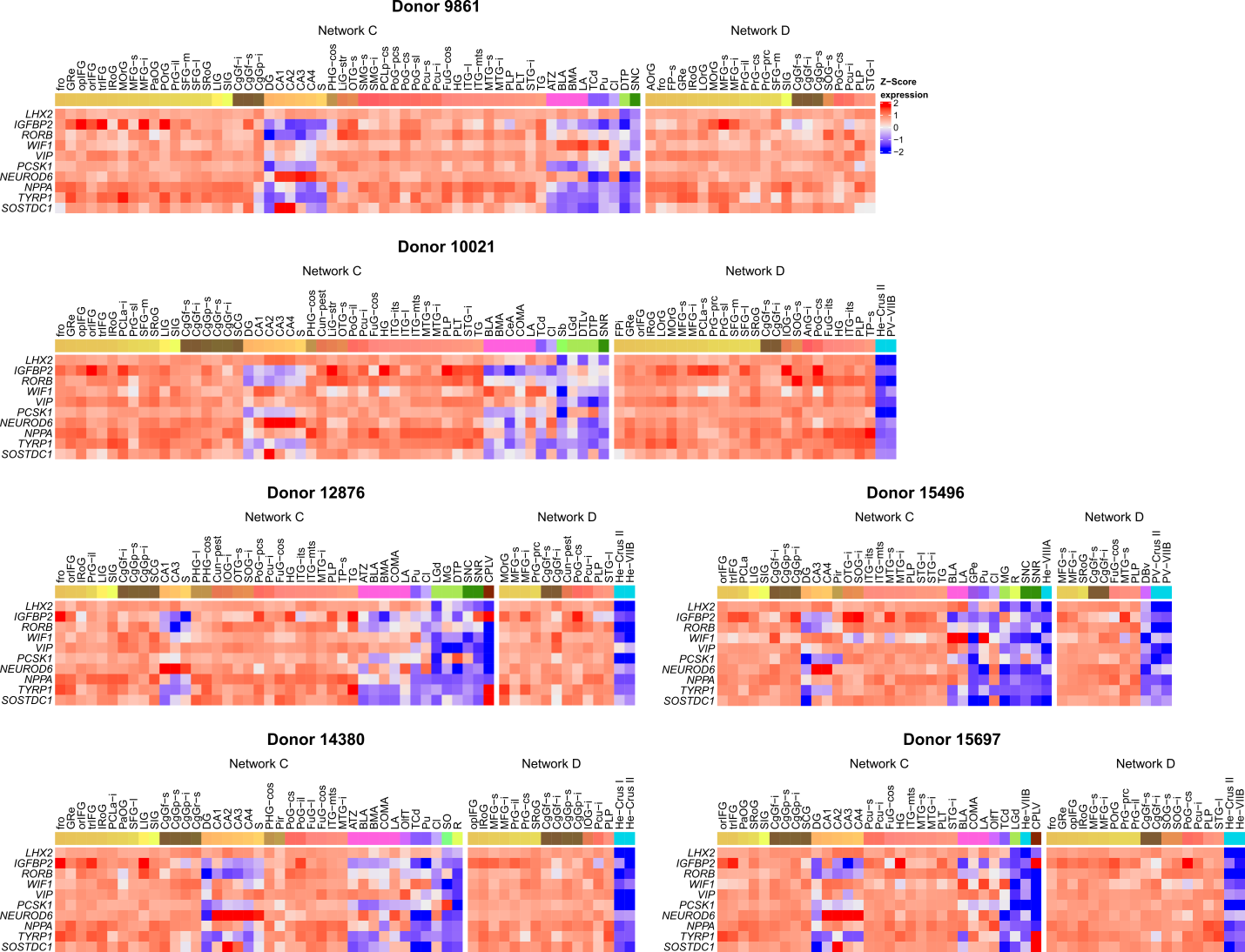

Supplementary Figure 2 Expression of differentially upregulated marker genes in the posterior cingulate network (C) and anterior cingulate network (D). Heatmaps of marker genes (rows) are shown for each one of the six donors in the Allen Human Brain Atlas and the color annotation for samples (columns). Expression was averaged across samples with the same acronym ignoring left and right hemisphere annotations. See Supplementary Table 7 for region-specific acronyms.

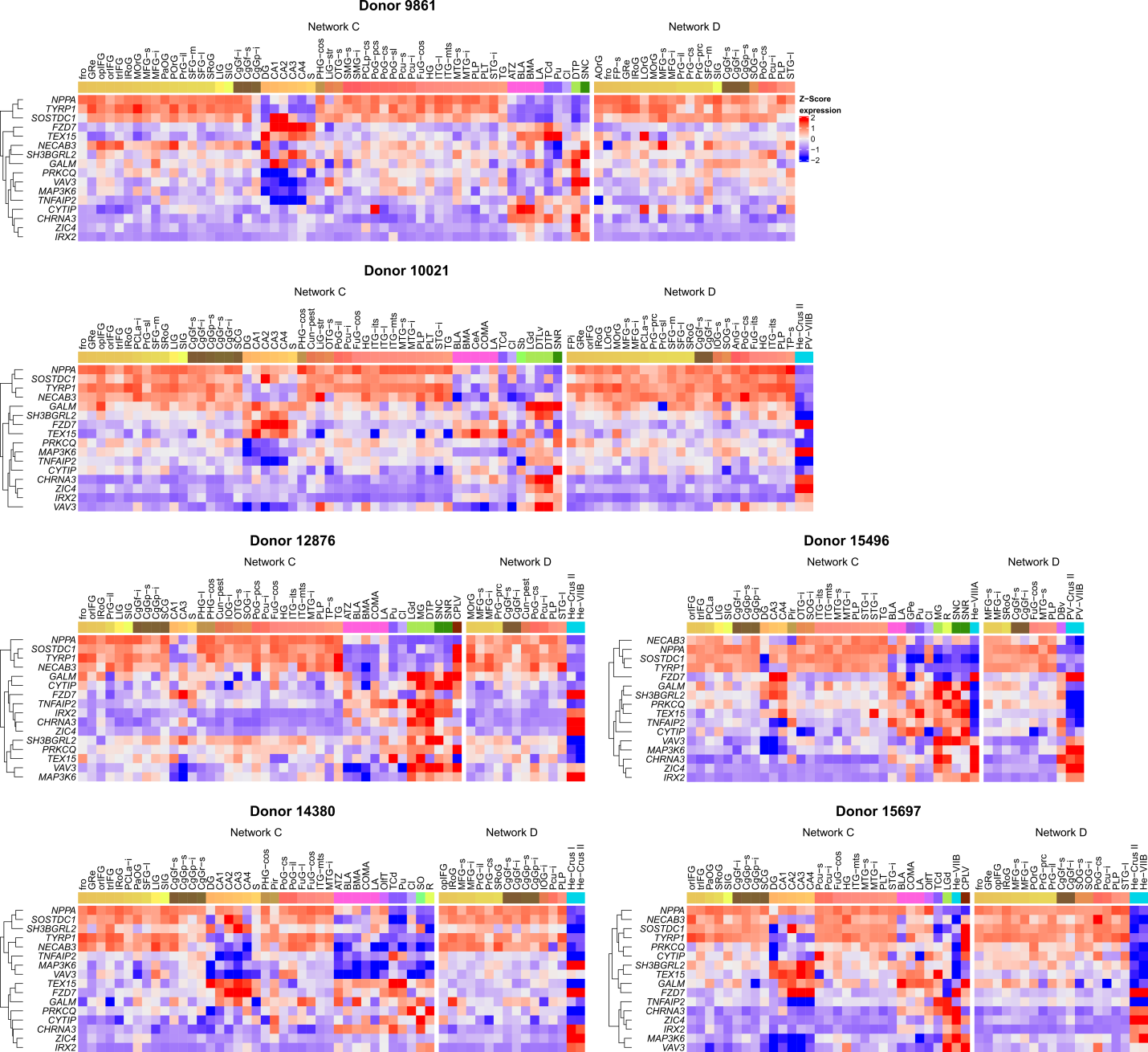

Supplementary Figure 3 Expression of thalamus cholinergic marker genes in the posterior cingulate network (C) and anterior cingulate network (D). Heatmaps are shown for each one of the six donors in the Allen Human Brain Atlas and the color annotation for samples (columns). Expression was averaged across samples with the same acronym ignoring left and right hemisphere annotations. Marker genes (rows) were clustered with complete linkage separately for each network. See Supplementary Table 7 for region-specific acronyms.
